# Maternal thyroid hormone increases neural cell diversity during zebrafish spinal cord neurodevelopment

**DOI:** 10.1101/2022.02.26.482108

**Authors:** Nádia Silva, Marco António Campinho

## Abstract

Maternally derived thyroid hormone (MT3) is a fundamental factor for vertebrate neurodevelopment. In humans, mutations on the T3 exclusive transporter monocarboxylic acid transporter 8 (MCT8) lead to the Allan-Herndon-Dudley syndrome (AHDS). Patients with AHDS present severe underdevelopment of the central nervous system with cognitive and locomotory consequences. Functional impairment of the zebrafish T3 exclusive membrane transporter MCT8 has been shown to phenocopy the symptoms observed in human patients with AHDS, thus providing an outstanding animal model to study this human condition. In this zebrafish model, MT3 acts as an integrator of different key developmental pathways during zebrafish neurodevelopment. Here we expand this knowledge by determining the developmental time of action of MT3 that occurs in very defined temporal intervals during zebrafish neurodevelopment. We have determined that MT3 is not involved in neural specification but is fundamental for developing particular neural progenitors and the consequent neural lineages that originate from them. Our data provide evidence that MT3 achieves this likely by modulation NOTCH signalling in a cell non-autonomous way. The findings show that MT3 expands the cell diversity output of neural progenitor cells, establishing a cellular background behind human AHDS and inherited limited CNS development.

## Introduction

Thyroid hormone (T3) metabolism has a high degree of regulation. However, the embryonic naïve system cannot endogenously produce the hormone during development, thus depending on a precise supply of maternal derived T3 essential for proper central nervous system (CNS) development. The outcomes of maternal thyroid hormone (MT3) deficiency in offspring are various and mainly depend on timing and severity of deficiency ^1^. It is well established that T3 influences neurodevelopment. Events influenced by MT3 during the foetal period comprise cellular proliferation, migration, and neuronal and glial cell differentiation (Williams, 2008). Although identified the genetic responses to T3 in specific cellular contexts ^2, 3^, and the phenotypic outcomes arising from inappropriate levels of TH supply are known ^1, 4^ the underlying cellular and developmental mechanisms are less understood. Furthermore, a key feature of T3 action during development is the strict windows of time where the hormone acts, which determines its action and biological outcome ^5, 6^. Nonetheless, the exact developmental times where MT3 action occurs during vertebrate neurodevelopment are unknown.

In humans, mutations in the exclusive T3 transporter, MCT8, are the cause behind the Allan-Herndon-Dudley Syndrome (AHDS) ^7, 8^. This syndrome, characterised by developmental delay, reduced myelination, intellectual disability, poor language and walking skills and hypotonia ^9^. The consequences of AHDS highlight the fundamental role of MT3 on vertebrate neurodevelopment. Although the pivotal role of MT3 on neurodevelopment is clear, there is still a significant knowledge gap on the genetic and cellular consequences of impaired T3 signalling. Furthermore, treatment of patients with T3 or its analogues cannot recover the neurological consequences of AHDS, highlighting the temporal specificity of T3 action during neurodevelopment ^10^.

Unlike in murine models ^11^, in zebrafish models of AHDS and thus MCT8 knockdown ^12–14^ it is possible to reproduce all the neurological consequences observed in human patients with AHDS. Evidence from previous zebrafish studies showed that several neural progenitors and neurons depend on MT3 for their development ^13, 15^. Furthermore, the spinal cord seems to be especially reliant on MT3 action for its normal development. In MCT8 morphant embryos, the spinal cord neurons dorsal and medial neurons are mostly lost or present abnormal morphology and positioning. In contrast, ventral neurons are favoured and increase their number in MCT8 morphant spinal cords ^13^. Transcriptomic analysis of MCT8 morphant embryos revealed that MT3 modulates several key developmental networks, like NOTCH, WNT and HH signalling, thus working as an integrative signal ^15^. Nonetheless, fundamental questions on the action of MT3 in zebrafish development and AHDS remain unanswered.In the present work we focused on spinal cord development and aimed to elucidate: 1) the developmental time window where MT3 action occurs, 2) the types of neural cell populations dependent on MT3 signalling and, 3) the cellular mechanisms by which MT3 acts.

In this study, we focused on the spinal cord and started to elucidate these fundamental questions on the action of MT3 in zebrafish development and AHDS. We show that MT3 signalling acts in well defined time windows from as early as 12hpf and most notably at 22-25hpf; MT3 signalling is required to develop of different neural cell types thus constituting an essential factor generating the cell diversity needed for appropriate central nervous system function and organisation. We provide further evidence suggesting that MT3 modulates neural progenitor lineage output via NOCTH signalling in a non-autonomous way.

## Results

### Timing of MTH action in zebrafish embryogenesis

To determine the developmental time window of MT3 action in zebrafish embryogenesis, we used the previously published transcriptome data at 25hpf ^15^. We selected genes regulated by MT3 at 25hpf involved in the early neural specification, NOTCH signalling pathway and neurogenesis (Fig. 1A).

**Figure 1.**
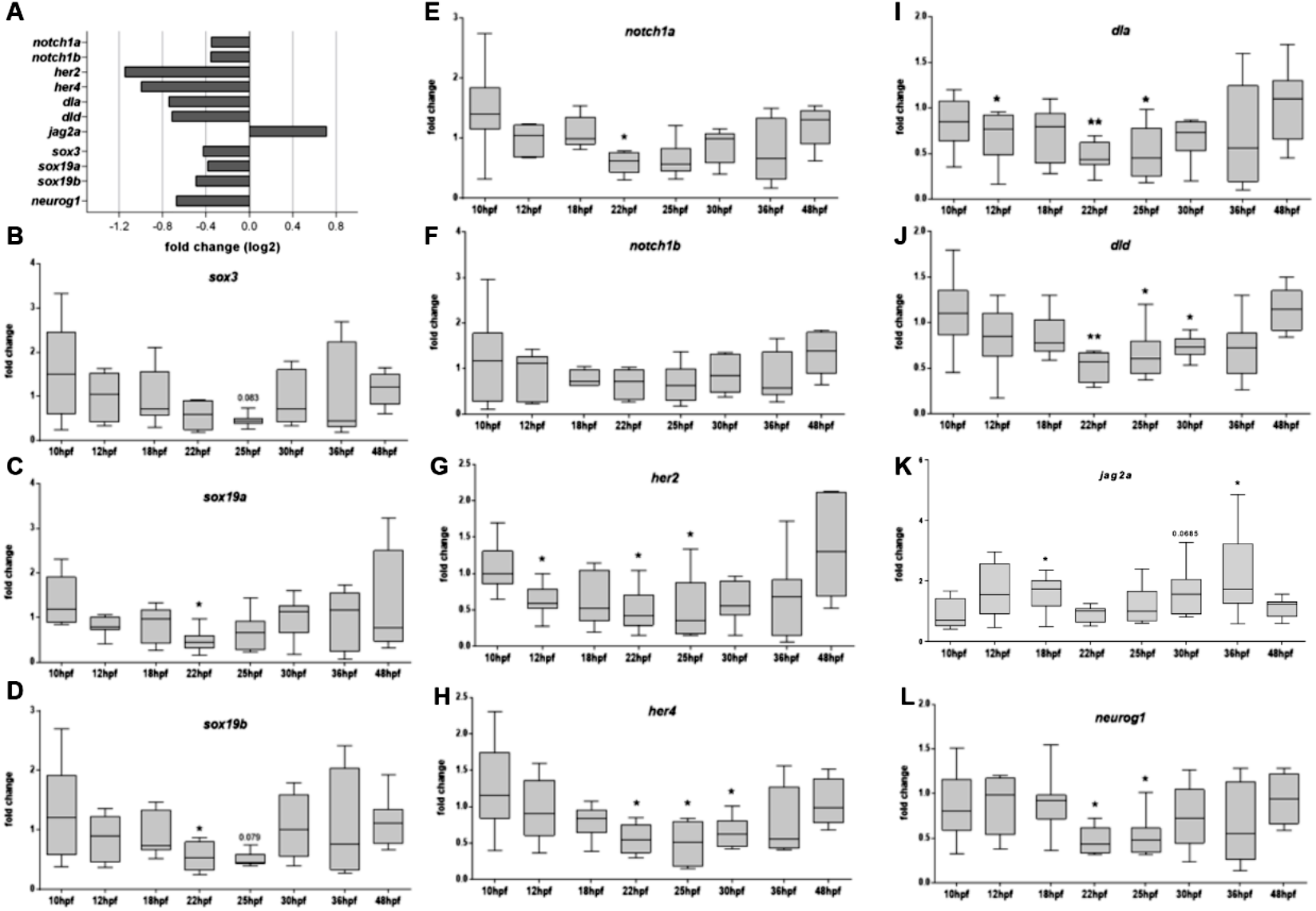
Expression of MT3-responsive genes reveals 22-31hpf as the developmental time more sensitive to MT3. (A) Quantification of genes of interest mapped to the neurogenesis cascade, differentially expressed between control and MCT8MO zebrafish, identified in RNA-Seq expressed as Log2 of fold change (n=7, p<0.01, FDR<0.0001) (NCBI –BioProjects: PRJNA381309). Box-and-whiskers plot of gene expression levels determined after RT-qPCR for *sox3* (B), *sox19a* (C), *sox19b* (D), *notch1a* (E), *notch1b* (F), *her2* (G), *her4* (H), *dla* (I), *dld* (J), *jag2a* (K) and *neurog1* (L). Data is represented as fold change of MCT8MO expression relative to the CTRLMO. Statistical significance was determined using a t-test: two-sample, assuming equal variances. N = 8 (* p<0.05; ** p<0.01).

Three SoxB1 genes (*sox3*, *sox19a* and *sox19b*) recognised for their role in the specification and development of the embryonic ectoderm into the neuroectoderm lineage ^16, 17^ were analysed. We aimed to determine if MT3 affects neural induction and contributes to the establishment of the neural plate by 10hpf. The candidate genes, *sox3*, *sox19a* and *sox19b*, were downregulated at 25hpf in the MCT8MO RNA-seq data (Fig. 1A), indicating a possible role MT3 at this stage in neural differentiation and maintenance of the neuroectodermal progenitor pool. Gene expression analysis by qPCR (Fig. 1B-D) showed that expression of these genes did not change in MCT8MO embryos during early neurodevelopment (10hpf-18hpf). The results suggest that MT3 does not play a role in maintaining B1 Sox gene expression during neural plate establishment and neural induction. Expression of *sox19a* and *sox19b* is significantly lower in MCT8MO embryos at 22hpf (t-test, p<0.05), while *sox3* and *sox19b* show a decreased expression also at 25hpf, in accordance with RNA-seq data ^15^, although this change does not reach statistical significance in the qPCR assay (t-test, p=0.083 and p=0,079, respectively).

Notch ligand-receptor combinations that coincide during development in zebrafish are essential for adequate brain development and cell diversity ^18^. To further understand how MT3 might be involved in neuronal cell fate determination and the timing of its action, we analysed the expression of Notch signalling components already known from previous transcriptomic data to have altered gene expression in MCT8MO embryos ^15^(Fig. 1A).

Gene expression analysis by qPCR revealed that the only significant difference between MCT8MO and CTRLMO zebrafish occurred for *notch1a* at 22hpf (Fig. 1E; t-test, p<0.05). Nonetheless, *notch1a* expression continues to be lower, although not statistically significant, in MCT8MO embryos until 36hpf (Fig. 1E). A similar trend occurs for the expression of *notch1b* receptor. However, the observed decrease in *notch1b* expression in MCT8MO embryos between 22-36hpf is not statistically significant (Fig. 1F, t-test, p>0.05).

The expression of notch direct targets *her2* (Fig. 1G) and *her4* (Fig. 1H), which are involved in the maintenance and proliferation of progenitor cell identity ^19–21^, decreases in specific time points. Expression of *her2* downregulates in MCT8MO embryos at 12, 22 and 25hpf (Fig. 1G, t-test, p<0.05), demonstrating an effect of MT3 on the Notch pathway during early neurogenesis. In zebrafish, *her4* is involved in the development of primary neurons under Notch 1 signalling ^21^. her4 downregulation at 22, 25 and 30hpf in the MCT8MO suggests the involvement of MT3 in regulating the development of some primary neurons (Fig. 1H, t-test, p<0.05).

Notch ligands *dla* and *dld*, which are expressed in differentiating neural cells, and are involved in the specification of progenitor pool size domains ^22^, showed a significant decrease in expression in MCT8MO embryos at 25hpf (Fig. 1I and J, respectively, t-test, p<0.05). The downregulation of *dla* is observable by 12hpf (Fig. 1I, t-test, p<0.05) during primary neurogenesis and also occurs at 22 and 25hpf (Fig. 1I, t-test, p<0.001 and p<0.05, respectively). The decrease in *dld* expression (Fig. 1J) only occurs later in neurogenesis at 22 (t-test, p<0.01), 25 (t-test, p<0.05) and 30hpf (t-test, p<0.05).

In contrast with *dla* and *dld*, the Notch ligand *jag2a* is upregulated MCT8MO embryos (Fig. K, t-test, p<0.05). The temporal pattern of expression determined by qPCR during development for *jag2a* was opposite to the delta ligands, *dla* and *dld* since it was upregulated at 18hpf (Fig. 1K, t-test, p<0.05) and again at 36hpf (t-test, p<0.05). Interestingly *jag1b*, another Notch ligand, was also upregulated, whereas *jag1a* was downregulated at 25hpf in the RNA-seq data (NCBI –BioProjects: PRJNA381309)^15^.

To further understand how MT3 is involved in neuron progenitor specification, we analysed the expression of *neurog1*, a pro-neural gene expressed by intermediate neuronal precursors and neuron committed cells (Fig. 1L). No differences in *neurog1* expression occur from 10-18hpf between CTRL and MCT8MO embryos suggesting MT3 is not involved in the differentiation of these cells. However, at 22 and 25hpf (Fig. 1L, t-test, p<0.05), *neurog1* expression decreased, suggesting a possible role for MT3 in the maintenance/differentiation of neuron progenitor populations from these stages of neurogenesis. After that, no differences in *neurog1* expression were found (Fig. 1L, t-test, p>0.05).

### Impaired MT3 action has a time-dependent effect on spinal cord neural development

The results from qPCR analysis of neural genes responding differentially to impaired MT3 action revealed that the hormone acts in a time-specific manner during zebrafish neurodevelopment (Fig. 1). We interrogated if these changes in gene expression are paralleled with changes in neurogenesis and gliogenesis. We focused our attention on the spinal cord since it provides a simplified version of neural development.

Immunostaining for Elav3 (HuC/D), which labels all zebrafish post-mitotic neurons from 15-48hpf, revealed a topological and different abundance of neurons in a time-dependent manner (Fig. 2) that parallels qPCR gene expression analysis of selected neural genes (Fig.1). Notably, in any developmental stage analysed, the distribution of neurons in the spinal cord is not similar between CTRMO and MCT8MO embryos (Fig. 2A). From as early as 15hpf, neurogenesis is impaired as can be seen by the decrease in HuC+ cells (Fig. 2B; t-test, p<0.05), the most affected spinal cord neuron population in MCT8MO embryos is medial, as can be observed in lateral and transversal sections (Fig. 2A). As development progresses, at 22hpf, there are fewer neurons in MCT8MO embryos (Fig. 2; t-test, p<0.05). All three regions of the spinal cord present different HuC staining profiles. In any axis, neuron distribution is different in control and MCT8MO embryos at 22hpf (Fig. 2A). That is especially evident in lateral view, where medial and ventral neurons seem to be particularly affected. By 25hpf, and although neuron numbers have recovered (Fig. 2B; t-test, p>0.05), the distribution of neurons is strikingly different from MCT8MO and control embryos (Fig. 2A). MCT8 morphant embryos neuron distribution at this stage is more compact (Fig. 2A transversal view) with an apparent accumulation of dorsally located neurons, some of which seem to be out of the spinal cord scaffold (Supplementary Figure 1). Neuron numbers decrease again at 36hpf (Fig. 2; t-test, p<0.0001). Additionally, the different distribution of the cells between control and MCT8MO embryos is exacerbated at 36hpf, where dorsal neurons seem to increase with a simultaneous decrease in medial and ventral neurons. By 48hpf, there is no difference in neuron number (Fig. 2; t-test, p>0.05), but the distribution of neurons in any view of the spinal cord is different in control and MCT8MO embryos. That is especially evident dorso-ventrally (Fig. 2A lateral and transversal views).

**Figure 2.**
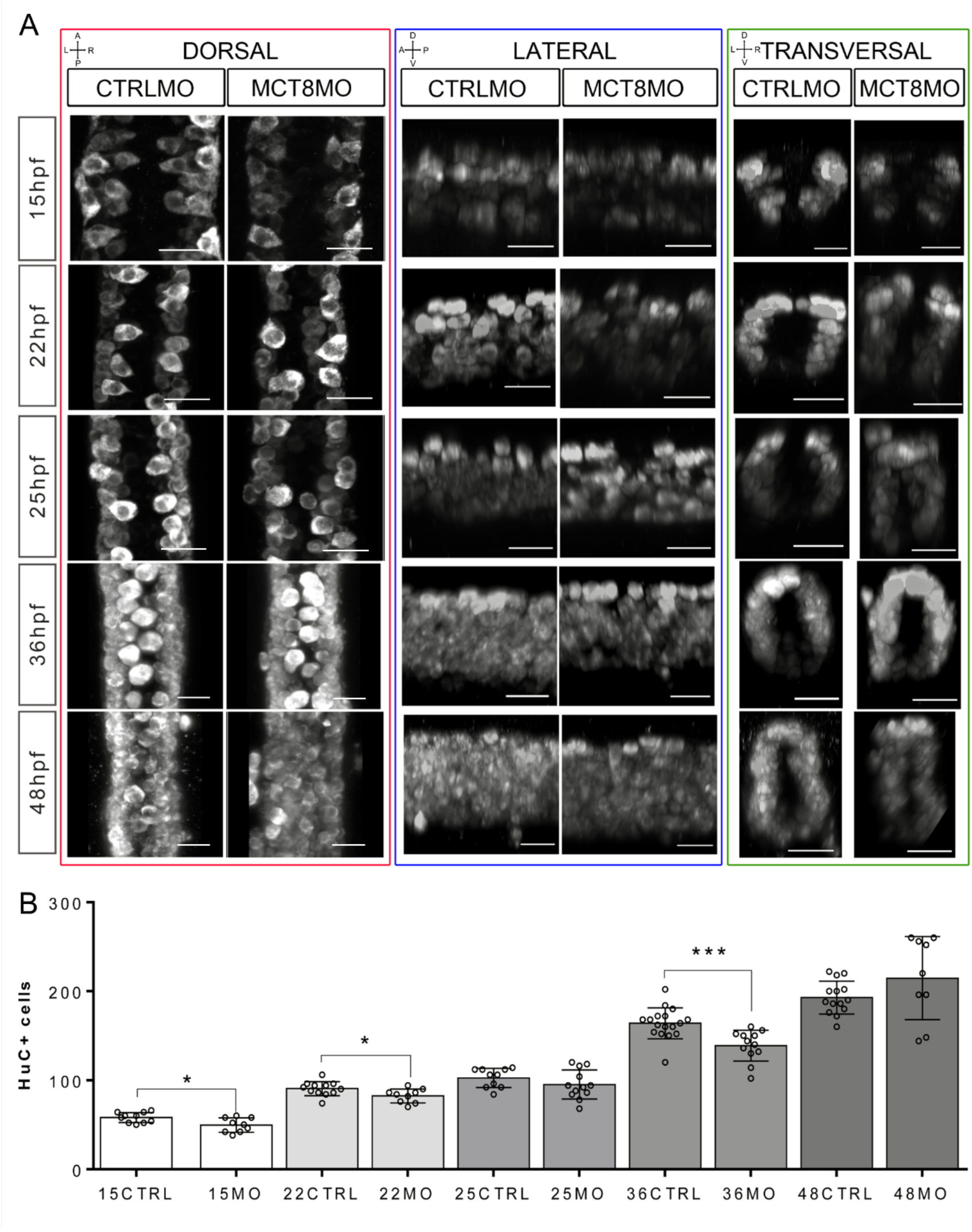
The number and distribution of neurons (HuC/D+) in the spinal cord of MCT8MO embryos is compromised at specific stages of development. (A) Representative maximum projection images of the pan-neuronal marker HuC/D immunostaining (white) in the spinal cord between somite 8-12. Comparison of the pattern of neuron distribution in the spinal cord between CTRLMO and MCT8MO embryos at different stages of development. (red highlight) Dorsal views, anterior spinal cord up. (blue highlight) Lateral view, anterior spinal cord right. (green highlight) Transversal view, dorsal spinal cord up. Scale bars represent 25 µm. (B) Quantification of the number of HuC/D single positive cells in a 2 myotome length of the spinal cord. n=9-17. CTRL (CTRLMO); MO (MCT8MO). Results are presented as mean±SD; Statistical significance determined by t-test: two-sample, assuming equal variances: *p<0.05; ***p<0.001.

These observations parallel the qPCR results of neurogenic related genes. Moreover, even though there is a recovery at the end of embryogenesis (48hpf), neuron distribution further suggests that specific neuron populations in control and MCT8MO embryos are necessarily different.

We also interrogated how spinal cord gliogenesis was affected by impaired MT3 signalling (Fig. 3). To this end, embryos were immunostained with an anti-GFAP serum (Sigma) and afterwards determined stained volume in similar trunk segments of the spinal cord. Likewise, even though we used a more restricted developmental stage range, we observed a time-dependent effect of impaired MT3 action on gliogenesis (Fig. 3). At 15hpf, there is a very significant decrease in GFAP staining in MCT8MO embryos (Fig. 3, t-test, p<0.001) with a very restricted GFAP signal (Fig. 3A). In contrast to CTRLMO embryos, in MCT8MO morphants, the ventral signal of GFAP at 15hpf was spread along the left-right axis of the spinal cord, whereas medial and dorsal staining was mostly lost (Fig. 3A, transversal). By 22hpf, the overall stain by GFAP was lower in MCT8MO than in CTRLMO embryos (Fig. 3, t-test, p<0.01). Notably, the topology of GFAP staining was different in control and MCT8MO embryos (Fig. 3). By this time, the GFAP signal in MCT8MO embryos increases in the lateral basal edge of the spinal cord, and little to no signal was found in the apical region.

**Figure 3.**
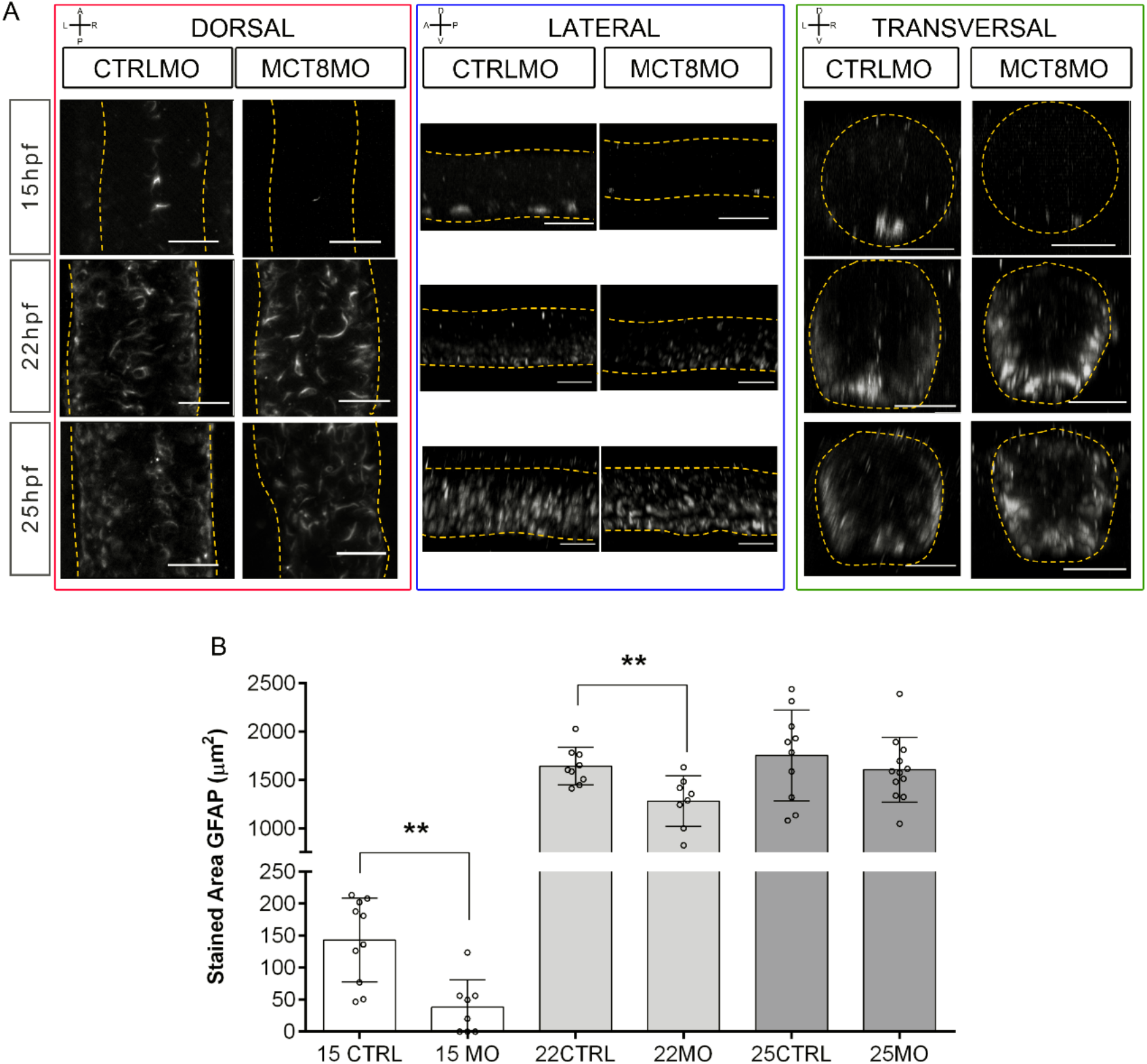
MCT8MO have altered glial cell development during early neurogenesis. (A) Representative maximum intensity projection images of the spinal cord between somite 8-12 after glial cell labelling with ZRF-1 immunostaining (white, labelling GFAP fibers). In control embryos at 15hpf glial cell fibers are organized in the developing ventral spinal cord; in MCT8MO embryos the development of these cells is delayed and only some scattered GFAP fibers are detected in the ventral-most neural tube. At 22hpf the neural tube is closed, and glial cells can be detected throughout the spinal cord of CTRLMO and MCT8MO embryos; at 25hpf patterning of glial cells is altered in MCT8MO embryos. Red box - dorsal views, anterior spinal cord up. Blue box - Lateral view, anterior spinal cord right. Green box - Transversal view dorsal spinal cord up. All scale bars represent 25 µm. Dashed yellow lines denote spinal cord boundaries. (B) Quantification of the area of GFAP staining in a 2 myotome length of the spinal cord. n=9-17. CTRL (CTRLMO); MO (MCT8MO). Results are presented as Mean±SD; Statistical significance determined by t-test: two-sample, assuming equal variances: **p<0.01.

By 25hpf, the stained volume of GFAP in the spinal cord is similar in both control and MCT8MO embryos (Fig. 3B, t-test, p>0.05). However, the signal distribution in any axis differs between the two experimental groups (Fig. 3A). Whereas GFAP staining in control embryos lined the basal edge of the spinal cord in MCT8MO embryos, GFAP staining was scattered throughout the basal-apical orientation of the spinal cord (Fig. 3, transversal). The observations argue that MT3 modulates gliogenesis, determining the position of glial cells and likely the cell diversity generated in this neural population.

To further dissect which cell populations in the spinal cord are affected by lack of MT3, we analysed the expression of genes involved in neural progenitor specification (*her2*, Fig. 4A), neuron committed progenitors (*neurog1*, Fig. 4B), radial glial cells (*fabp7a*, Fig. 4C), astrocyte-like cells (*slc1a2b*, Fig. 4D), oligodendrocytes (*olig2*, Fig. 4E) and motoneurons (nkx6.1, Fig. 4F).

**Figure 4.**
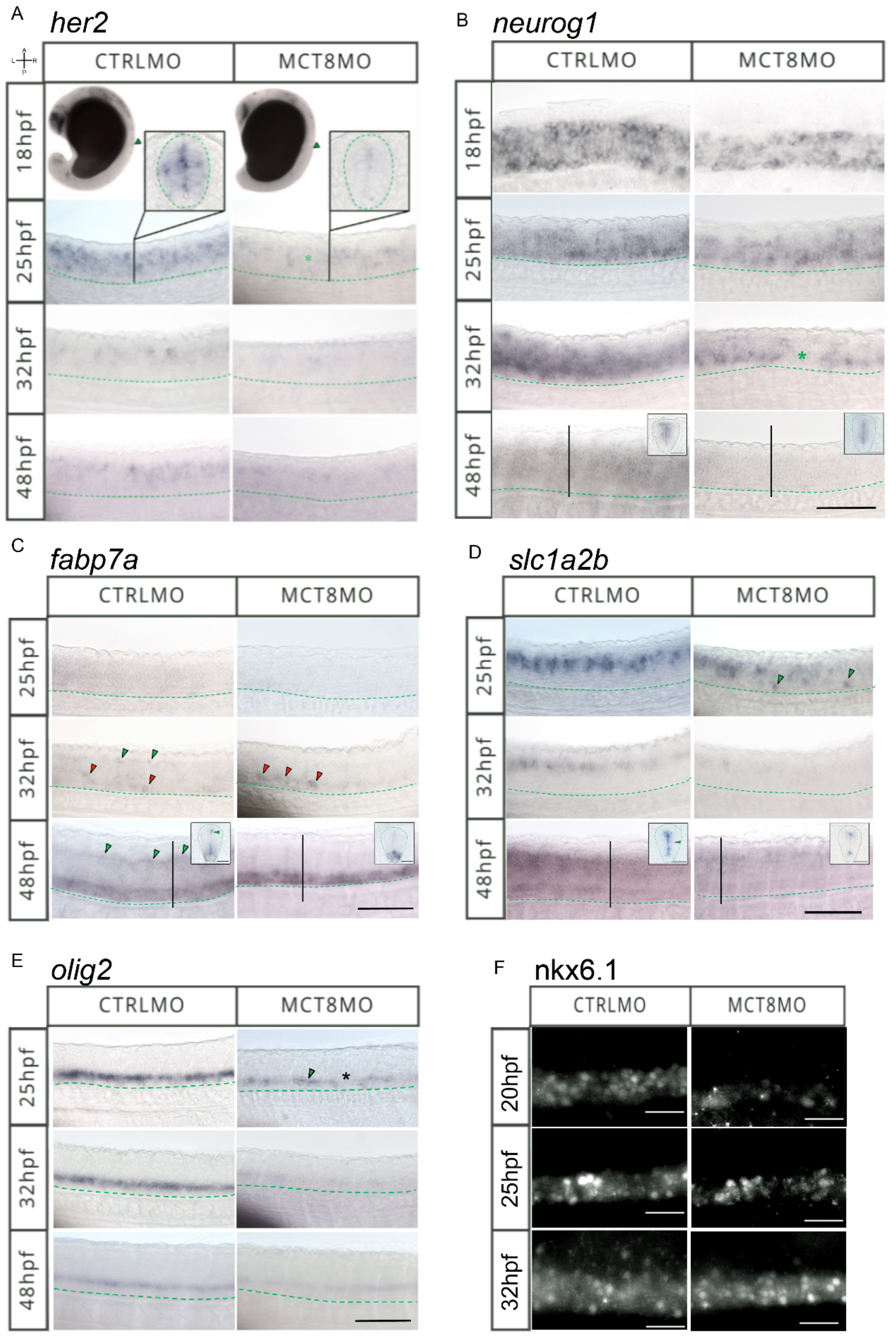
MTH is necessary for the correct development and positioning of different neural cells in the spinal cord of zebrafish. (A) Expression pattern of *her2* detected by WISH in whole embryo at 18hpf and the spinal cord at 25, 32 and 48hpf. The green asterisk at 25hpf represents a spinal cord region were *her2* expression is absent in MCT8MO embryos. Insets are vibratome transverse sections of 25hpf CTRLMO and MCT8MO were the difference in expression pattern between CTRL and MCT8MO is more noticeable. (B) *neurogenin-1* (*neurog-1*) expression pattern in the spinal cord using WISH at 18hpf (dorsal view), 25, 32 and 48hpf (lateral views). Inserts at 48hpf are vibratome transverse sections of the CTRL and MCT8MO embryos, where expression of neurog1 in the sc dorsal domain of MCT8MO embryos is compromised. The green asterisk in 32hpf MCT8MO highlights a spinal cord area were *neurog1* expression is absent. (C) *fabp7a* spinal cord expression pattern after WISH at 25, 32 and 48hpf. At 32hpf in CTRLMO embryos the green arrowheads indicate the dorsal *fabp7a*+ cells, which are lost in MCT8MO embryos. Red arrowheads indicate *fabp7a* staining in the ventral domain of the spinal cord, which is increased in MCT8MO embryos. At 48hpf, green arrowheads indicate the presence of *fabp7a*+ cells in the dorsal domain of CTRLMO spinal cord while they are less evident in MCT8MO. This is more noticeable in the insets of vibratome transverse sections of the region indicated by a black vertical line in CTRL and MCT8MO 48hpf embryos. (D) *slc1a2b* spinal cord expression pattern after WISH at 25, 32 and 48hpf in CTRL and MCT8MO embryos. At 25hpf green arrowheads indicate the position of misplaced *slc1a2b*+ cells in the ventral spinal cord of MCT8MO embryos compared to CTRLMO. At 32hpf expression of *slc1a2b* is less abundant in MCT8MO. CTRLMO embryos at 48hpf display a broad expression region of *slc1a2b* surrounding the neurocoelom, while MCT8MO embryos display a reduced signal for *slc1a2b* in cells at the most ventral and dorsal regions of the neurocoelom. Insets at 48hpf are vibratome transverse sections of the region indicated by the vertical black line in CTRLMO and MCT8MO. The green arrowhead in CTRLMO indicates the spinal cord area in which *slc1a2b* expression is less abundant in MCT8MO embryos. (E) *olig2* spinal cord expression pattern after WISH at 25, 32 and 48hpf in CTRLMO and MCT8MO embryos. In CTRMO embryos from 25 to 48hpf *olig2*+ cells were present in the most ventral region of the spinal cord. The green arrowhead indicates the position of *olig2*+ cell clusters that were present and asterisk denotes the absence of cells in regions of the spinal cord in MCT8MO embryos at 25hpf, when compared to CTRLMO. Overall MCT8MO embryos presented less intense staining for *olig2* with apparently less *olig2*+ cells present, this was observed in all stages. (F) Nkx6.1 immunofluorescence detection in the spinal cord at 20, 25 and 32hpf. In all stages analysed Nkx6.1 protein showed an apparently decrease presence in MCT8MO, when compared to CTRLMO. In all images the green dashed line represents the ventral limit of the spinal cord. The images present a lateral view of the spinal cord (unless specified) between somite 8-13. Rostral is to the left in all images. Scale bar represents 50 µm . In all insets the green dashed line in represents the outermost boundary of the spinal cord and the scale bar represents 20 µm. A minimum of 10 individuals/conditions/gene were analysed.

*her2*+ neural progenitors are lost from as early as 18hpf, and dorsal populations are most affected (Fig. 4A). Cells expressing *her2* becomes more spaced, suggesting that some but not all *her2*+ progenitors are more susceptible to impaired MT3 signalling than others (Fig. 4A).

A similar situation is observed for *neurog1*+ neuron committed progenitors (Fig. 4B). At 18hpf, there are significantly fewer *neurog1*+ cells (Fig. 4B). By 25hpf dorsal *neurog1*+ cells are mostly lost in MCT8MO embryos (Fig. 4B). This pattern continues at 32hpf, but additionally, it is observed gaps in *neurog1*+ cells in the spinal cord (green asterisk in Fig. 4B). That observation suggests that specific *neurog1*+ progenitors at specific spinal cord locations are more dependent on MT3 than others. By 48hpf, *neurog1*+ progenitors in MCT8MO embryos are restricted to the more ventricular region of the spinal cord that surrounds the neurocoelom (inserts in Fig. 4B).

At 25hpf *fabp7a*+ radial glial cells seem to be highly dependent on MT3 for their development, given that almost no cells are present in MCT8MO embryos (Fig. 4C). By 32hpf, some *fabp7a*+ cells are found, but these are mostly ventral (red arrowheads in Fig. 4C), whereas dorsal cells are lost (green arrowheads in CTRLMO embryos in Fig. 4C). That accentuates at 48hpf where no dorsal *fabp7a*+ cells (green arrowheads in CRTLMO embryos in Fig 4 C) are found in MCT8MO embryos but only ventral located cells. Moreover, the ventral expression field of *fabp7a*+ cells in MCT8MO embryos is more extensive and present a different spatial distribution (inserts in Fig. 4C).

Astrocyte-like cells expressing *slc1a2b* are also affected in MCT8MO embryos (Fig. 4D). Already by 25hpf, there is a decrease in expression of *slc1a2b* in the spinal cord of MCT8MO embryos (Fig. 4D) with a less dense row of cells present. Accompanying these expression changes and in a dorsal location, the development of ventral located *slc1a2b*+ cells in MCT8MO embryos (green arrowheads in Fig. 4D) is also observed not present in control embryos. By 32hpf, there is a general decrease of *slc1a2b*+ cells in MCT8MO embryos spinal cord (Fig. 4D). By 48hpf, the general decrease of *slc1a2b*+ cells observed in MCT8MO embryos is accompanied by a restriction of the expression field that by this time confined to the most dorsal and ventral regions of the spinal cord canal (inserts in Fig. 4D).

In MCT8MO embryos, *olig2*+ cells in the spinal cord decrease from as early as 25hpf (Fig. 4E). This reduction of *olig2*+ cells seems to be higher at 32hpf but slightly recovers by 48hpf (Fig. 4E). Nonetheless, *olig2*+ staining is always lower in MCT8MO than in control embryos (Fig. 4E), suggesting loss of some cells.

We also labelled by immunohistochemistry nkx6.1 expressing cells in the spinal cord of experimental embryos (Fig. 4F). From as early as 20hpf, nkx6.1+ cells strongly decrease in MCT8MO embryos. That continues up until 25hpf, but at 32hpf, there seems to be an expansion of nkx6.1 cells in MCT8MO embryos (Fig. 4F). Nonetheless, this partial recovery in MCT8MO embryos cells with a high signal for nkx6.1 is more abundant than in control embryos (Fig. 4F). Moreover, while in control embryos, a medial to a dorsal expansion of nkx6.1 cells occurs in MCT8MO embryos these are mostly concentrated in a ventral position (fig. 4F), strongly suggesting that the identity of these nkx6.1+ cells is not identical in control and MCT8MO embryos.

### MT3 is essential for a subset of neural progenitor cells to survive and proliferate

The previous results suggest that MT3 is involved in specifying distinct neural populations. The decrease in *her2, neurog1,* and *fabp7a* expression in MCT8 morphants suggest that the function of MT3 in the generation of cell diversity in the zebrafish spinal cord arises already at the progenitor level, either by restricting the fate of daughter cells or restricting the diversity within the progenitor pool itself.

All components of T3 cellular signalling (i.e. *mct8*, *thraa* and *thrab*) are already present in the zebrafish neuro-epithelium from as early as 12hpf and widely expressed in the spinal cord up until 48hpf (Supplementary Figure 2). At 25hpf, there are several MT3 sensitive (*mct8+*) *her2+* neural progenitors in a scattered pattern and dorso-ventrally more frequent in the ventral half (Fig 5A). Likewise, both *thraa* and *thrab* are co-expressed with *her2* in the spinal cord (Fig 5A). The co-expression pattern with *her2* of the receptors is different from the one with *mct8* but also between themselves. Whereas *thraa* is mostly co-expressed with *her2* dorsally (arrow) and continuously expressed anteroposteriorly, *thrab/her2* co-expression appears, anterior-posteriorly, in bands separated by regions of no co-expression (Fig 5A). These *thrab/her2* co-expression bands spawn the dorsal-ventral axis but are more frequent in the medial region of the spinal cord. In MCT8MO embryos, co-expression of *her2* with the receptors is not lost, but it decreases and presents different distributions (Fig 5A). In mct8 morphant embryos, *thraa/her2* co-expression becomes more ventral (arrowhead) and medial, even though some dorsal co-expression (Fig 5A). *thrab/her2* co-expression loses the anterior-posterior pattern of defined bands and becomes continuous and more restricted to the medial region of the spinal cord (Fig 5A). These findings indicate the existence of at least six her2+ neural populations dependent at some point of their spinal cord development of MT3 signalling components: MT3+/thraa/+thrab+, MT3+/thraa+/thrab-, MT3+/thraa-/thrab+, MT3-/thraa+/thrab+, MT3-/thraa+/thrab- and MT3-/thraa-/thrab+.

**Figure5.**
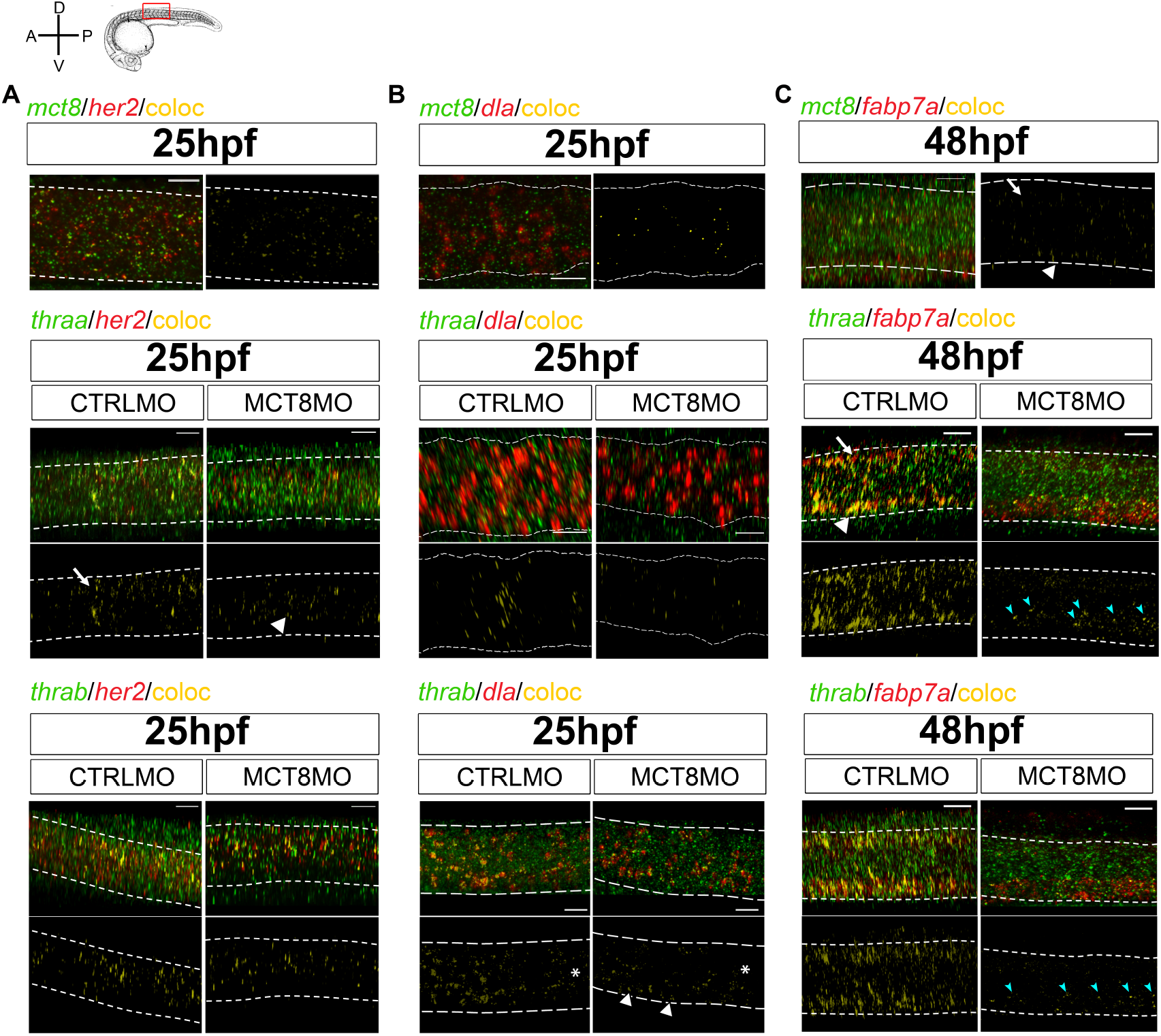
MT3 is directly involved in development of discrete her2, dla and fabp7a+ cells. Colocalization of zebrafish *thraa, thrab* and *mct8* with *her2* (A) and *dla* (B) expressing cells at 25 and *fabp7a* (C) at 48hpf after double whole-mount in situ hybridization. *thraa, thrab* and *mct8* (green); *her2, dla* and *fabp7a* (red) and colocalization (yellow). *thraa*/*her2* colocalization in CTRLMO (arrow) embryos is increased in the apical spinal cord, while in MCT8MO *thraa*/*her2* colocalization has a more medial distribution in the spinal cord (arrowhead). *thrab*/*dla* colocalization is less predominant in MCT8MO embryos, asterisks denote decreased colocalization along the anterior-posterior axis of the spinal cord. Arrowheads indicate the increased colocalization of *thrab/dla* (+) in cells of the ventral spinal cord in MCT8MO embryos. *mct8/fabp7a* colocalization in CTRLMO occurs scattered through the spinal cord with an arrow indicating colocalization in the dorsal spinal cord and arrowhead colocalization in the ventral spinal cord. Colocalization of *fabp7a* with *thraa* and *thrab* in MCT8MO embryos is more restricted to the ventrally located *fabp7a+* cells (blue arrowheads). All images depic a spinal cord section between somite 8-12; anterior is left and dorsal up. White dashed lines show the boundary of the spinal cord. A minimum of 3 individuals/conditions was analysed. Scale bar represents 20µm.

At 25hpf, MT3 sensitive cells (*mct8+*) *dla+* cells restrict to the medial region of the spinal cord. No *mct8/dla* co-expressing cells are detected in the spinal cord’s most dorsal and ventral regions (Fig 5B). At this time, spinal cord *thraa/dla* co-expression has a very defined anterior-posterior expression pattern in bands that spawns dorso-ventrally but is more frequent medially (Fig 5B). In contrast, *thrab/dla* co-expression has an anterior-posterior decrease in frequency (asterisks) but uniformly distributed dorso-ventrally and with the presence of large clusters (Fig 5B). At 25hpf, mct8 morphant embryos co-expression of *dla* with *thraa* or *thrab* is severely decreased, and its distribution changed compared to control siblings (Fig 5B). In these embryos, *thraa/dla* co-localisation loses the pattern found in control siblings and is scattered with some larger clusters found in discrete dorsal, medial and ventral regions of the spinal cord (Fig 5B). In turn, *thrab/dla* co-localisation still presents a decreased anterior-posterior expression (asterisk) but is almost lost dorsally and medially (Fig 5B). Although decreased, compared to control embryos, the co-expression of *thrab/dla* is more frequent ventrally (arrowheads in Fig.5B). These observations indicate that only a small population of *dla+* progenitors cells depends on MT3. Since most *thraa+/dla+* co-expression is lost in MCT8 morphant embryos, in the MT3 sensitive, *mct8+*/*dla+* population cells, *thraa* is likely the most important receptor mediating the genomic hormone action. In MCT8 morphants, a similar situation occurs for *thrab+/dla+* co-expression, although not so widespread. Moreover, comparing the patterns of co-expression between control and MCT8 morphant embryos indicates that: i) in most dorsal and ventral regions of the spinal cord, *thrab/dla+* cells are likely mostly irresponsive to MT3 and; ii) most MT3 sensitive *thrab+/dla+* cells have a medial localisation (Fig 5B).

We further look at *fabp7a* co-localisation with *mct8*, *thraa* and *thrab* at 48hpf, a time of extreme sensitivity of RGC to MT3 (Fig. 4C). In control embryos, MT3(*mct8+/fabp7a+*) sensitive RGC cells are localised primarily on the most dorsal (arrow) and ventral (arrowhead) regions of the spinal cord (Fig 5C). As well, most *fabp7a*+ cells co-express *thraa+* and/or *thrab+* (Fig 5C), indicating that RGCs are highly dependent on MT3 regulated transcription. That becomes even more evident when we look to MCT8 morphant embryos, where there is a very drastic decrease in the frequency of *fabp7* co-expression with *thraa* and *thrab* (Fig 5C). More for *thrab* than for *thraa*, co-expression with *fabp7a* is almost entirely lost dorsally (Fig 5C). As most of *the fabp7a* signal is lost, it indicates that most dorsal RCG cells depend on *fabp7a* to develop. Ventrally, a different scenario is found in MCT8 morphants. Although most co-expression of *fabp7a* with *thraa* and *thrab* is lost, *fabp7a* expression increases (Fig 5C). Notably, the remaining co-expression fields of *fabp7a* with either *thraa* or *thrab* are in clusters on the dorsal portion of the most ventral third of the spinal cord (cyan arrows in Fig 5C). Nonetheless, the superimposition of the two co-expression fields does not retire the possibility that in some *fabp7a+* cells, MT3 action occurs via both receptors (Fig 5C). Together these observations indicate that: i) dorsal developing *fabp7a+* RGC rely more on MT3 to differentiate than ventral RGC; ii) ventrally MT3 action might be more important in RGC fate decisions and diversity (specialisation) generation than general RGC *fabp7a+* development; iii) most *fabp7a+* RGC are dependent on MT3 genomic action, but a small portion of RGC depend on thyroid receptor aporeceptor function to develop.

To further understand how MT3 acts on spinal cord *her2+* and *dla+* neural progenitors development, we carried out assays to understand if these cells stop proliferating or undergo apoptosis in the absence of the hormone (Figs. 6 and 7). We analysed control and MCT8 morphant embryos between 18hpf and 25hpf, a time where qPCR analysis showed that *her2* and *dla* expression was more responsive to MT3. In general, cell proliferation (as measured from mitotic index labelling PH3) is decreased by ∼50% on average at 18 to 25hpf (Supplementary Figure 3A, p<0.0001). Albeit lower in MCT8 morphants, at 25hpf, the two experimental groups have a similar number of mitotic cells (Supplementary Figure 3A). On the other hand, in all developmental stages analysed, there is increased apoptosis in MCT8 morphants twice as much as in control embryos (Supplementary Figure 3B, Dunn p<0.001). As previous reported ^13^, this increase in apoptosis is specific to the lack of MT3 and cannot be rescued by p53 signalling abrogation, thus indicating a specific effect of impaired MT3.

**Figure 6.**
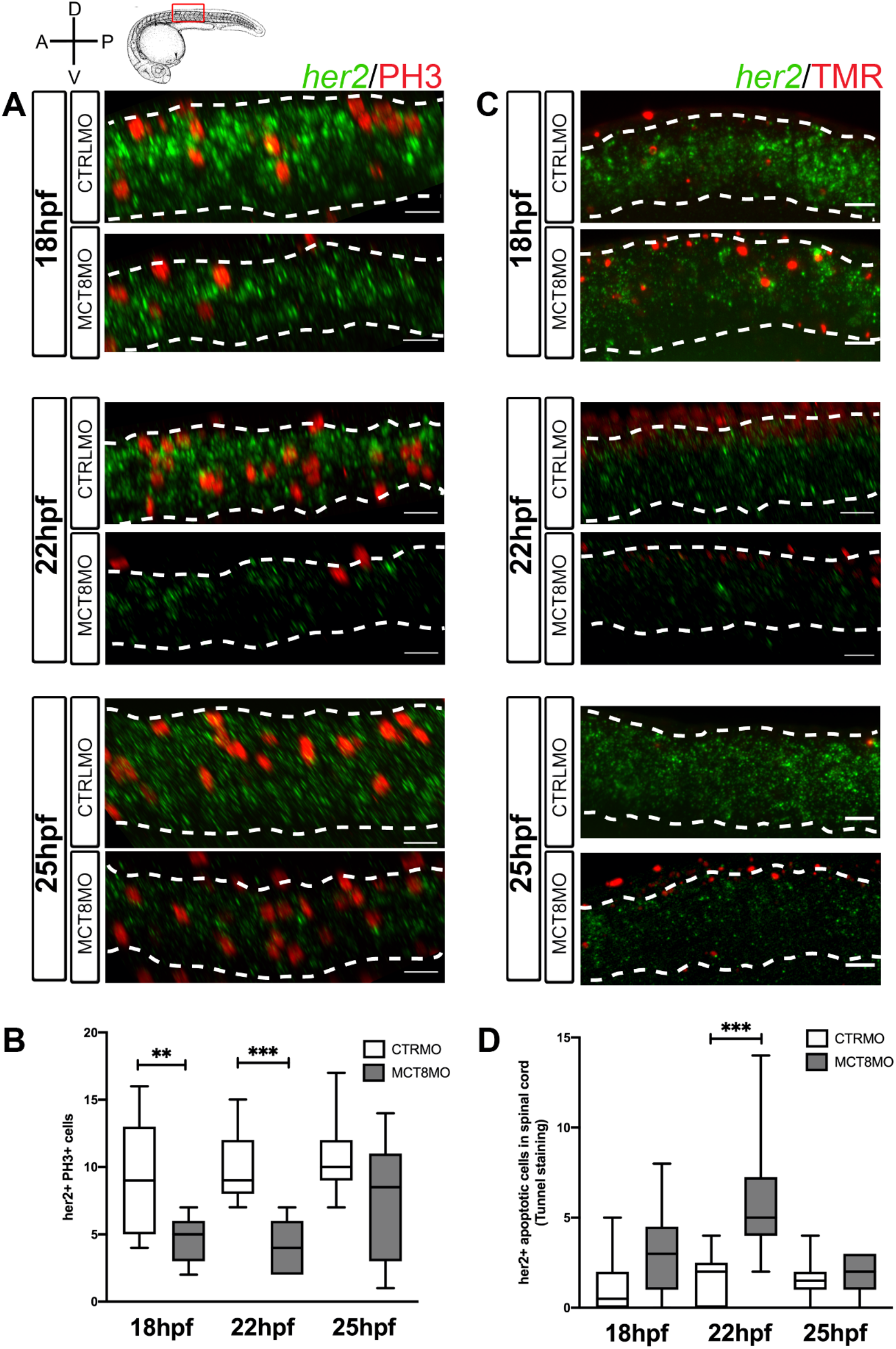
Impaired MT3 signalling decreases *her2+* neural progenitor cells undergoing mitosis in the spinal cord. (A) Analysis of *her2* expression by fluorescent *in situ* hybridization (green) and mitotic cells (phosphohistone 3 antibody; red) in control (CTRLMO) and MT3 impaired (MCT8MO) embryos. (B) Box-and-whiskers plot depicting quantification of the number of *her2*(+) mitotic cells (her2+/PH3+) in the spinal cord at 18, 22 and 25hpf. (C) Analysis of *her2* expression by fluorescent *in situ* hybridization (green) and colocalization with apoptotic cell detected using a TUNNEL assay (red). The images represent a lateral view of the spinal cord between somite 8-12, rostral is to the left in all images; scale bar is 50 µm. (D) Box-and-whiskers plot depicting the quantification of the number of *her2*(+) apoptotic cells in the spinal cord. Colocalisation quantification was made in 2 myotome volume within this region of the spinal cord. n=10-15. Statistical significance determined by t-test: two-sample, assuming equal variances. **p<0.01; *** p<0.001.

**Figure 7.**
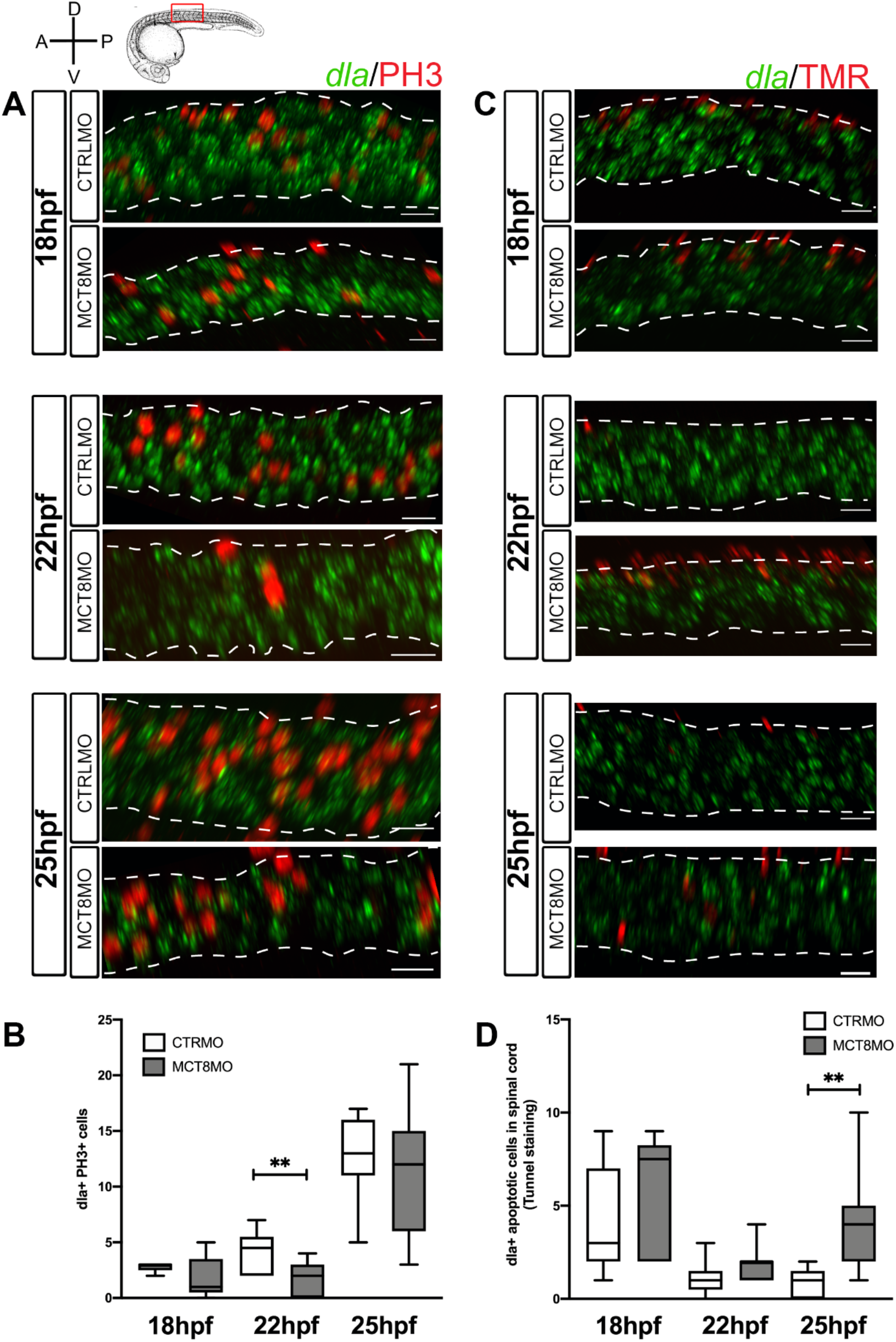
Impaired MT3 signalling affects proliferation and apoptosis of *dla+* spinal cord cells in a time restricted manner. (A) Analysis of *dla* expression by fluorescent *in situ* hybridization (green) and colocalization with cell mitosis (phosphohistone 3 immunostaining, red). The images present a lateral view of the spinal cord between somite 8-12, rostral is to the left and dorsal up in all images. Scale bar represents 50 µm. (B) – Box-and-whiskers plot of quantification of the number of *dla*(+) mitotic cells in the spinal cord at 18, 22 and 25hpf in control and MCT8MO injected embryos. (C) Analysis of *dla* expression by fluorescent *in situ* hybridization (green) and colocalization with apoptotic cell detected using a TUNNEL assay (red). (D) Box-and-whiskers plot of quantification of the number of *dla*(+) apoptotic cells in the spinal cord at 18, 22 and 25hpf in control and MCT8MO injected embryos. Countings were made in 2 myotome volume sections within this region of the spinal cord (n=10-15). Statistical significance determined by t-test: two-sample, assuming equal variances. **p<0.01.

At 18 and 22hpf, *her2*+ mitotic cells presented a decrease of ∼50% in MCT8 morphant embryos compared to control (Fig. 6A and B, p<0.01). At 25hpf, there are no differences between the two groups. These results parallel the obtained for general PH3 staining and suggest that about one-quarter of *her2+* mitotic cells depend on MT3 to proliferate (Fig. 6B and Supplementary Figure 3A). Loss of *her2+* mitotic cells in MCT8 morphants occurs more frequently in medium and ventral regions of the spinal cord at 18 and 22hpf (Fig. 6A). By 25hpf, there is an evident increase in *her2*+ mitotic cells in these regions of the spinal cord in MCT8 morphants (Fig. 6A).

Apoptosis of *her2+* cells in MCT8 morphants is only higher at 22hpf (p<0.001) but not at 18 and 25hpf (Fig. 6C and D). The divergence of *her2+* apoptotic cells from general spinal cord apoptosis indicates that only a small sub-set of *her2+* arising at 22hpf are likely dependent on MT3 to develop. Irrespective of any experimental group, apoptotic *her2+* cells are more frequent dorsally, especially at 22 and 25hpf (Fig. 6C). Together these observations indicate that from 18 until 22hpf, about one-quarter of *her2+* progenitors depend on MT3 to survive. MT3-dependent survival of *her2+* cells is restricted in space and developmental time interval around 22hpf (Fig. 6). Together, this evidence argues that the major role of MT3 on *her2+* progenitors is likely involved in cell fate decisions and cellular diversity generation.

*dla+* cells proliferation is not dependent on MT3 at 18 and 25hpf but only at 22hpf (Fig. 7A and B, p<0.01). At this time, only one-sixth of *dla+* cells are proliferating, and of these, only half seem to be dependent on MT3 (Fig. 7B). Notably, *dla+* proliferating cells do not follow the same frequency as the one observed for general proliferation in the spinal cord for both control and mct8 morphant embryos (Fig. 7A, B and Fig. Supplementary Figure 3A). In control embryos, a large proportion of *dla*+ proliferating cells occurs ventrally, and these are mostly lost in MCT8 morphants (Fig. 7A). That indicates that at 22hpf, *dla+* MT3-dependent proliferating cells are mostly ventrally localised (Fig. 7A).

In contrast to cell proliferation, *dla+* apoptotic cells in MCT8 morphants are only higher at 25hpf (Fig. 7C and D, p<0.01), accounting for twice as much as those found in control embryos. Moreover, *dla+* apoptotic cells in control embryos only represent about 20% of all apoptotic cells in the spinal cord at 25hpf, thus suggesting that in MCT8 morphant embryos, apoptotic *dla+* cells might represent a different *dla+* population than the one found in control siblings (Fig. 7D and Supplementary Figure 3B). Notably, *dla+* cell death does not follow the same distribution for overall spinal cord apoptosis (Fig. 7D and Supplementary Figure 3B). In both control and MCT8 embryos, most *dla+* cells are found in the most dorsal region of the spinal cord at 18 and 22hpf (Fig. 7C). In contrast, to control embryos, at 25hpf in mct8 morphants, *dla*+ apoptotic cells locate mainly in the medial and ventral regions of the spinal cord. The results indicate that only a small subset of *dla*+ cells depends on MT3 for proliferation and survival. Moreover, this dependence seems restricted to dorsally located cells and well-defined developmental times (Fig. 7).

The previous evidence further supports that NOTCH signalling mediates MT3 action in zebrafish spinal cord neural progenitor cells. Furthermore, our evidence support that the dependence of NOTCH signalling on MT3 is highest between 18-30hpf. To further understand if this action of MT3 can be cell-autonomously rescued by activated NOTCH signalling, we injected 1-cell stage embryos with 0.8pmol of CTRL or MCT8MO. Afterwards, between 16-32 cell stage we injected a DRA1 or DRA2 blastomeres in each morpholino injected group with either GFP mRNA or NICD+GFP mRNA, carried out live imaging of the spinal cord between 23-26hpf (Fig. 8A) and quantified symmetric and asymmetric GFP+ cells divisions (Fig. 8C-F). As criteria, we classified symmetric dividing cells if the cell division plane occurred from 0°<30° in respect to the basal side of the spinal cord or asymmetric divisions if these occur with an angle of ≥30° (Fig. 8B)^23^. There were no differences in overall cell division of GFP expressing cells between any of the experimental groups (Fig. 8C). In comparison to control embryos, there were differences in the proportion of symmetric/asymmetric divisions with other experimental groups (Fig. 8D; χ^2^, p≤0.05). Nonetheless, symmetric divisions occurred less frequently in NICD+CTRLMO, MCT8MO and NICD+MCT8 experimental groups; these were not statistically significant from the CTRLMO (Fig. 8E, t-test p>0.05). In contrast, asymmetric divisions in MCT8MO and NICD+MCT8MO experimental groups were significantly less frequent when compared to the CTRLMO group (Fig. 8F, One-way ANOVA p<0.01, Sidak, p<0.05) but not the NICD+CTRLMO or between themselves. These results argue that NOTCH overexpression cannot rescue the lack of MT3 signalling in these progenitor cells in a cell-autonomously manner.

**Figure 8.**
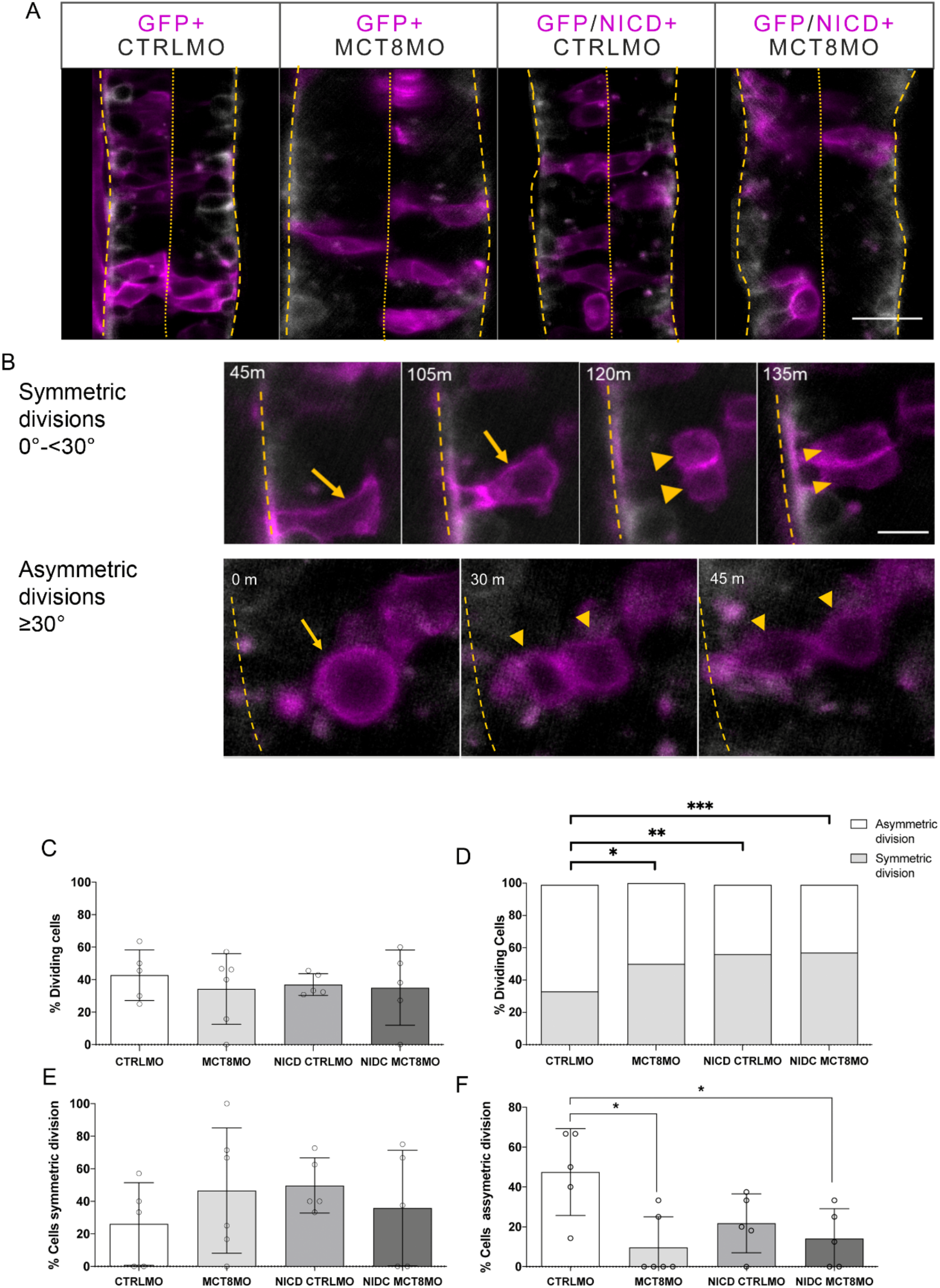
NICD overexpression cannot rescue in a cell-autonoumous manner the effects of impaired MT3 singalling on assymetric cell division in the spinal cord. (A) Representative images of spinal cord of experimental Tg(elav3:LY-mCherry) embryos at 23hpf at the start of imaging. Neurons (mCherry) are shown in white. Embryos were injected at one cell stage with either CTRLMO or MCT8MO followed by injected at the 16 cell stage in one blastomere with gfp mRNA only or nicd and gfp mRNA. Cells overexpressing NICD (and GFP) are labelled in magenta. Dorsal views of single slices between somite 8-15 are shown, anterior spinal cord up. Scale bar represents 50 µm. (B) Upper panel: Detail of symmetric division originating 2 morphologically similar GFP+ cells. The yellow arrow denotes a cell undergoing symmetric mitosis, the yellow arrowheads indicate daughter cells. Lower panel: Representative images of asymmetric division. Dividing cell (yellow dashed line) originates two GFP+ daughter cells (yellow line). The scale bars in B represent 20 µm. In all panels the spinal cord basal limit is indicated by a dashed yellow line and the pial limit by dotted yellow lines.. C-F: (C) Percentage of analysed cells GFP+ cells that underwent division. (D) Distribution of symmetric and asymmetric divisions on dividing cells observed from 23-26hpf. χ^2^ analysis showed differences in the distribution of the number of cell undergoing symmetric or asymmetric divisions amongst experimental groups and CTRLMO (p<0.05). (E) Percentage of cell undergoing symmetric division. (F) Percentage of cells undergoing asymmetric division. The results in C, D, F are presented as the mean±SD, results in D depict ratio of cell division type in all GFP+ dividing cells analyzed. N = 5-6 individuals per group (number of cells evaluated by group: CTRLMO/GFP=56; MCT8MO/GFP=77; CTRLMO/NICD/GFP=93; MCT8MO/NICD/GFP = 46); Statistical significance in C, E, F was determined by a one-way ANOVA followed by a Holm-Sidak’s multiple comparison post hoc analysis, *p<0.05.

## Discussion

The irreversible neural effects of pathologies arising from the early embryonic deficit of MT3 cannot be rescued by subsequent treatment with thyroid hormones or analogues. That is well reflected in human AHDS patients where post-natal T3 or T3 analogue treatment cannot rescue the neurologic consequences in these patients ^9, 24–26^. This constitutes evidence of the existence of a critical time window for T3 action during embryonic neurodevelopment. The zebrafish AHDS model used in the present work provides the first evidence of this critical developmental time of MT3 on vertebrate neurodevelopment. Our data support MT3 time-restricted action in zebrafish neurodevelopment. In this respect, 18-30hpf (∼pharyngeal stage) is found to be, based on T3-responsive gene expression, the most dependent on MT3. The data argue that MT3 is not required for neuroectoderm induction but generates cell diversity during neurogenesis and gliogenesis. That is likely achieved by MT3-regulation of particular neural progenitors’ developmental output (i.e. fate decisions) reflecting neuronal and glial cell populations. Our analysis strongly supports that besides a time-dependent effect, MT3 action on target cells is also dependent on tissue/cellular context. Our data support that in zebrafish, as in mammalian systems, T3 is involved in the differentiation and proliferation of a wide variety of cell types and that MT3 action on cell proliferation depends on the cell identity, developmental state and cellular context ^27^. However, at 12hpf, there is already a differential expression of notch ligand *dla* and *her2*, pointing out that a subset of neural progenitor cells also responds to MT3 early in neurodevelopment. The identity of these early MT3-responsive cells and their progeny remains to be confirmed.

We demonstrate that MT3 is essential for the proliferation and survival of both neural stem cells and committed neuron and glial progenitors. That argues that MT3-regulation of neural diversity is achieved by modulating the output from various progenitor cells. Impaired MT3 action on zebrafish neurodevelopment affects mostly dorsal her2+ neural stem cell, neurog1+ intermediate neuron progenitors, *fabp7a+* and *pax6a+*^15^ radial glial progenitors and *olig2+* motoneuron and oligodendrocyte progenitors. Cells arising from these progenitors like *slc1a2+* astrocytes, nkx6.1a+ motoneurons and *gad1b+* inhibitory interneurons^15^ are also restricted in their dorsal domains in mct8 morphants. In several animal models, it has been described T3 action over neuron development and survival. In chicken, mct8 knockdown leads to impaired optic tectum development, depletion of neuroprogenitors and impaired neurogenesis with reduced neuron numbers and diversity ^28^. In in vitro mammalian cells, T3 is directly involved in the development of granule neurons by, on the one hand affecting the survival and differentiation of these cells ^29^ but also by preventing their apoptosis^30^. In mammalian neurodevelopment with impaired maternal thyroid hormone signalling, either by embryos carrying thyroid hormone receptors with mutations giving resistance to T3 ^31–34^, congenital hypothyroid athyroid pax8 mutants or in double-knockdown mct8/oatp1c1 embryos^35, 36^, gave rise to similar cellular effects to the ones observed in zebrafish mct8 morphant embryos (^13, 15^, present study). In the developing mct8/octp1c1 double-KO embryonic mice cortex^36^, *Hr* and *gad67* (respectively, homologs of zebrafish *her2* and *gad1b*) expressing cells are mostly lost dorsally, strongly suggesting that in all vertebrates studied so far, MT3 constitute an important factor in dorsal specification of neuron cell identities most notably inhibitory neuron development. In rat embryos, T3 deficiency decreases the proliferation and delays the maturation of the precursors of cerebellar GABAergic interneurons, with effects on the number of mature GABAergic neurons and GABAergic terminals ^37, 38^. Notably, in mct8 morphants, it is observed a decrease in dorsal spinal cord neurons while is observed an increase in ventral motoneurons ^13^. Given the similarity of the observed locomotive phenotype in mct8 morphants ^12, 13^ and human AHDS patients ^25, 39, 40^ it is very likely that the cellular basis of impaired locomotion in human patients is related to increase in excitatory neurons and depletion of GABAnergic interneurons. Above all, a key observation in zebrafish mct8 morphants is the apparent recovery of spinal cord neuron numbers at 25 and 48hpf. However, the topology and morphology of these neurons and the neural circuits developed in the spinal cord are not identical to control morphants. That indicates that in the loss of MT3-dependent neuron development, other neuron types assume their positions/location, thus anticipating a compensatory mechanism. A similar compensatory mechanism was observed in Xenopus neurodevelopment. Here impaired NOTCH signalling leads to delayed neurogenesis but is later compensated at the expense of impaired cell diversity ^41^. In fact, in human AHDS patients, microcephaly is rarely observed ^25^ strongly suggesting that impaired development of some neuron types leads to overgrowth from other types. Indeed, in chicken embryonic retinal development, mct8 knockdown leads to a shift towards increased blue cones at the expense of green/red cones ^42^, further arguing that MT3 is involved in all vertebrates in generating neuron cell diversity and the adequate balance between all neuron types in order to be able to develop a fully functional central nervous system.

In wild-type zebrafish embryos, co-expression of *her2* with *mct8* resembles mostly *her2* co-expression with *thraa*, suggesting that in *her2+* progenitors, effectuation of MT3 signalling is thraa driven. Indeed, in mct8 morphants, *thraa* co-expression with *her2* is mostly lost in dorsal spinal cord cells, whereas it is mostly maintained ventrally. That argues that *her2+* dorsal NSC populations are dependent on MT3 action via thraa, whereas ventral populations rely on thraa unliganded aporeceptor function to differentiate. A similar but not predominant situation seems to occur with thrab since medial spinal cord co-expression with *her2+* is mostly maintained in mct8 morphants but lost dorsally. From our analysis, the loss of dorsal *her2+* MT3-dependent progenitors is likely due to apoptosis since TUNNEL staining strongly co-localises with *her2+* cell in mct8 morphants. A similar situation is found in the embryonic mouse cortex where impaired MT3 leads to decreased cell cycle length and apoptosis of progenitors cells ^3, 43, 44^. Moreover, in cultured rat pituitary tumour granule cells, T3-induced cell proliferation is mediated by changes in G1 cyclin/cyclin-dependent kinase levels and activity ^45^. However, from our analysis, one cannot discard that decreased *her2+* MT3 dependent cells diminished their numbers after reduced proliferation due to preconscious differentiation and exit from the cell cycle. Previous transcriptomic analysis in zebrafish mct8 morphants show a steep decrease in expression of cell-cycle genes ^15^ that further strengths this possibility. Another possibility is that the lack of MT3 leads these progenitors into a state of senescence. This has been observed to be the case in adult *mct8*/*octp1c1* double-KO mice mutants^46^.

Our data also suggest that different progenitor populations respond to MT3 differently. Although co-expression of *her2* and *dla* was previously established by single-cell analysis in wild-type zebrafish embryos^47^, the effect of MT3 absence on proliferation and apoptosis of *her2* and *dla* expressing cells is unequal. *dla* is expressed in neural precursors and transiently in post-mitotic neurons at 11.5hpf ^48^. The increased cell death of progenitor cells, especially at an early stage of neurogenesis, can be a contributing factor in reducing progenitor pools leading to compromised neurogenesis. That is the case for oligodendrocyte progenitor populations in zebrafish spinal cord development ^49^. In the case of *dla+* cells, regulation by MT3 seems to depend on two different mechanisms where thraa and thrab have different roles. *thraa+*/*dla+* cells present a vertical band like pattern with high co-localisation in the medial region of the spinal cord and are highly dependent on MT3. In the case of *thrab+*/*dla+* cells, only dorsal located cells are dependent on MT3. From this observation, it is clear that at least three different *dla+* populations exist: one that is dependent on MT3 and relies on thraa; one dorsal *thrab+* population that is dependent on MT3 and; a ventral population that is positive to *thrab* but likely irresponsive to MT3. Nonetheless, it is not yet possible to determine the identity and the progeny arising from these different *dla+* cell populations.

Interestingly, our data suggest that zebrafish developing *fabp7a+* radial glial cells are highly dependent on MT3 and that both trhaa and thrab are fundamental for the response of these cells to MT3. Additionally, our data also points out that glial cells are dependent on the precise timing of MT3 signalling, as observed by the decrease in *fabp7a+* and *slc1a2b+* staining. A similar situation occurs in the developing mouse hippocampus and cerebellum, wherein hypothyroid embryos GFAP expression was markedly reduced in a time-dependent manner ^50^. Again, from our data, dorsal *fabp7a+* RGC co-localise with *thraa* and *thrab* and these are almost entirely lost in mct8 morphants. In contrast, although decreased, ventral co-localisation of *fabp7a+* RGC with either *thraa* or *thrab* is still maintained in mct8 morphants leading to an expansion of *fabp7a+* RGC cell expansion in the ventral domain. Interestingly, in mct8 morphants, *pax6a+* is also lost dorsally but less so ventrally^15^ further arguing for a differential dorsal-ventral role of MT3 in RGCs development. From these observations, it is clear that MT3 is involved in the specification of different *fabp7a+* RGCs in the spinal cord, which is then reflected in the restricted development of *slc1a2b+* astrocyte like cells in mct8 morphant embryos. Collectively, our data indicate that MT3 is essential to establish the correct combination of glial cell types that allow the development of the adequate cytoarchitecture of the spinal cord. The finding further supports that in zebrafish mct8 morphants, it is observed that neurons develop outside of the dorsal spinal cord, a region where the greatest loss of RGCs is observed.

The developmental genetic mechanisms underlying MT3 control of development are still poorly understood. Notwithstanding, our previous findings indicate that MT3 regulates zebrafish neurodevelopment by modulating important genetic signalling pathways, most notably, WNT, SHH and NOTCH pathways ^15^. In zebrafish neurodevelopment, the NOTCH pathway seems to be especially responsive to MT3 signalling as major system components respond in a time-dependent manner to the hormone (present study). Importantly, NOTCH plays a fundamental role in the regulation of animal neurodevelopment (revised in ^51^), most notably by lateral inhibition, where it promotes cell fate specification of neural progenitors and daughter cells. However, the only examples of T3 control of NOTCH pathway come from studies in mouse ^52^ and Xenopus ^53^ post-natal intestinal development. In mouse and Xenopus intestinal post-natal development, T3 regulates the action of the NOTCH pathway in intestinal progenitor cells hence functioning as a cell fate determinant. In these cases, several components of the NOTCH pathway, including receptors and ligands, are regulated by T3 in a time and cell-context dependent manner ^52^ ^53^. Our findings point to a similar mode of action of MT3 on the NOTCH pathway in regulating neural progenitor proliferation, survival and developmental output during zebrafish neurodevelopment (discussed above).

Using live imaging, we show that impaired MT3 signalling decreases asymmetric divisions during zebrafish spinal cord development while not affecting symmetric divisions. In the developing nervous system, symmetric divisions are associated with progenitor pool amplification or terminal differentiation of progenitors. In contrast, asymmetric divisions are related to the acquisition of new cell fates by daughter cells or asymmetric terminal differentiation giving rise to different daughter cells and, in this way, increasing in cell diversity ^54^. Together with our previous data, this evidence strongly supports that MT3 is necessary for developing particular cells fates and generating cell diversity during neurodevelopment. Furthermore, mosaic overexpression of NICD (and thus cell-autonomous activation of the NOTCH pathway) cannot rescue in a cell-autonomous manner the consequences of impaired MT3 signalling in neural progenitor cells. That observation further argues that MT3 likely functions in neurodevelopment as an integrative signal that allows for balanced NOTCH signalling that gives rise to the different neural cell types in a time and cell context-dependent manner and that cannot be rescued in a cell-autonomous manner. That is further supported by the fact that MT3 regulates NOTCH receptors, delta and jagged ligands expression. The observation that delta and jagged ligands expression is regulated in opposing manners by MT3 (^15^, present study) suggests that the hormone functions as a balance and integrator that enables the appropriate input from NOTCH ligands and the developmental outcome that arises from that. This is of extreme significance given that new studies indicate that NOTCH ligand dynamics are fundamental for mice multipotent pancreatic progenitor cells output and the fate of daughter cells arising from the division of these progenitors ^55^. Such an integrative function of MT3 in neurodevelopment supports the observations that both excess and impaired hormone signalling have profound effects on central nervous system development and function. Nonetheless, new studies are necessary to dissect further MT3 role in NOTCH signalling modulation and neural progenitor output in zebrafish neurodevelopment.

The implications of present findings for the comprehension of ADHS, and the development of putative therapies, are immense. It has been suggested that the pathogenesis associated with MCT8 deficiency arises from an impaired T3 transport across the blood-brain-barrier ^56^. Here we show that the effect over neural cell progenitors occurs prior to blood-brain-barrier development in zebrafish. They show that the MT3 entering through cellular mct8 of CNS-residing cells regulates their development.

Although several treatments for AHDS syndrome have been undertaken, and in some cases, the peripheral thyrotoxicosis was ameliorated, the underlying cause for this neurological phenotype could not be rescued ^10^. The timing of T3 action is essential. In the first trials of treatment for neurological cretinism by iodisation, researchers realised treatment was only efficient if it occurred before pregnancy or during the early stages of foetal development (reviewed in ^57^). Although the phenotype of these two pathologies is not identical, since they have diverse aetiologies, they both arise from the fact that T3 did not reach the target cells during a particular developmental time window. Overall, our data support that: i) the restricted temporal action of MT3 is critical for vertebrate neurodevelopment; ii) key vertebrate neuron types dependent on MT3 for development are inhibitory neurons, whereas glial cells (RGCs and astrocytes) seems to be generally dependent on MT3 for development; iii) In zebrafish mct8 morphant embryos and likely human AHDS patients the overall neurodevelopmental effects of MT3 impairment arise from both lack of direct action of MT3 on target gene transcription and as well as relief of gene expression repression by unliganded thyroid receptors; iv) in both cases the likely cause behind this impaired development is decreased differentiated neural cell diversity due to loss of lineage-committed progenitors.

Given this evidence, two non-mutually exclusive hypotheses arise to explain how MT3 regulates vertebrate neurodevelopment: 1) MT3 acts in neural progenitors in order to allow particular cellular states that enable the generation of the full potential cell fates arising from these progenitors and/or 2) MT3 acts by allowing final differentiation and survival of neural progenitors committed to a given cell fate generation.

## Materials and methods

### Zebrafish husbandry and spawning

Adult wild-type (AB strain) zebrafish were maintained in standard conditions in the CCMAR zebrafish fish facility at the University of Algarve (Portugal). Fish husbandry was carried out according to the EU Directive on protecting animals used for scientific purposes (2010/63/EU). Adult fish were kept at a 14 h/10 h light/dark cycle and 28°C. Breeding stock feeding twice a day with granulated food (Tetra granules, Germany) and once with Artemia sp. nauplii. One female and one male zebrafish were isolated in mating tanks the night before egg collection. The separator was removed when the lights turned on in the morning.

### Morpholino injection

Upon spawning, embryos were immediately collected and microinjected at the 1-2-cell stage with 1nL of morpholino solution containing either 0.8pmol CTRLMO or MCT8MO as described in (Campinho et al., 2014). Embryos were randomly distributed into plastic plates containing E3 medium (5 mM NaCl, 0.17 mM KCl, 0.33 mM CaCl, 0.33 mM MgSO4) and incubated until sampling time at 28.5°C (Sanyo, Germany) under 12h:12h light:dark cycles.

### Analysis of gene expression

The staging was after observed developmental landmarks in control embryos (Kimmel et al., 1995). Eight independent biological replicates (pools) of 20 embryos were sampled at the first five-time points (10, 12, 18, 22, 25 hpf). Eight biological replicates (pool) of 15 embryos were sampled from 30, 36 and 48hpf embryos; Embryos were manually dechorionated, snap-frozen in liquid nitrogen and stored at -80°C. According to the instructions manual, total RNA from the embryos was extracted with an OMEGA Total RNA extraction kit I (Omega Biotek, USA). To remove genomic DNA, total RNA was then treated with an Ambion Turbo DNA-free kit (Life Sciences, USA) accordingly to the manufacturer’s instructions. The concentration of total RNA was determined after the NanoDrop ND-1000 spectrophotometer (NanoDrop Technologies Inc., USA). RNA integrity was determined after visualised on an agarose gel stained with SYBR Green nucleic acid gel stain (ThermoFisher Scientific).

500ng of purified total RNA was reverse transcribed to complementary DNA (cDNA) using RevertAid First Strand cDNA Synthesis and Random Hexamer Primers (Thermo Fisher Scientific, USA) according to the manufacturer’s instructions. Synthetized cDNA was diluted 1/5 in ultrapure water and stored at -20°C.

The quantification method used with the RT-QPCR method was the absolute quantification method, which determines the number of mRNA copies in the sample from a standard curve. For that, a standard curve for each gene of interest was prepared. Primers were designed using Primer 3 Plus using RNA-seq data^15^. Table 1 provides primer sequences and amplicon size for each gene included in the analysis.

**Table 1.**
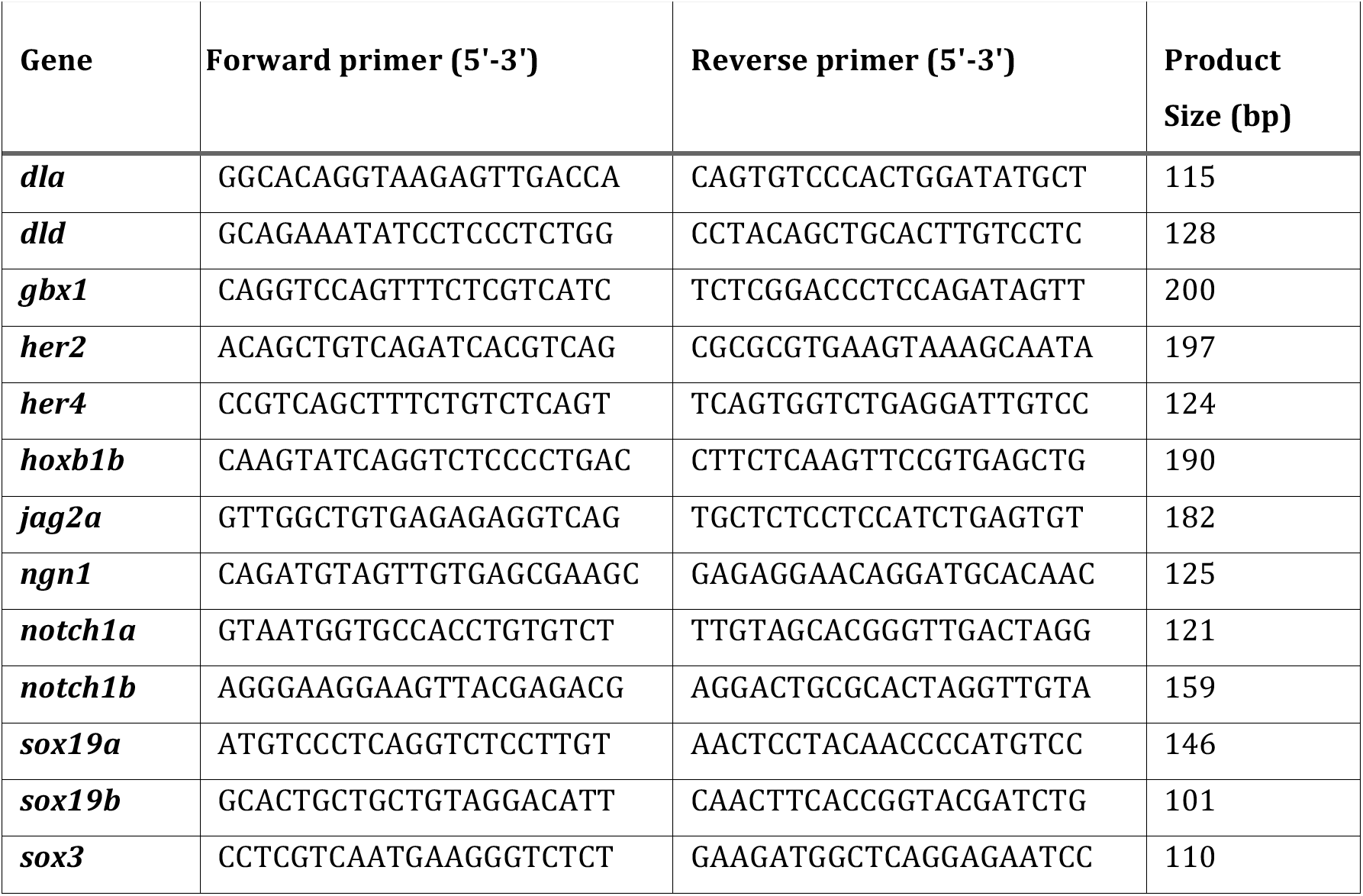
List of primers for qPCR analysis.

The gene’s target sequence was amplified by PCR, purified (EZNA Gel Extraction Kit, Omega Biotek), quantified (NanoDrop Technologies Inc., USA) and sequenced by Dye-termination to confirm their identity.

Quantitative real-time PCR (qRT-PCR) was performed in a CFX-384 well (Biorad). qPCR was carried out in a 384-well format with 6 µL of total volume. Final concentrations of PCR mix consisted of 1X SensiFASTTM SYBR, No-ROX Kit (Bioline, USA), 150nM forward primer, 150nM reverse primer, 1 µL cDNA (1/5). The PCR amplification protocol was 95 °C for 3 min, and 44 cycles of 95 °C for 10 sec and 60 °C for 15 sec. The amplification protocol included a denaturation step from 60 to 95°C, 5 sec in 0,5°C increment, to obtain a standard melting curve to confirm the production of a single amplicon and absence of primer dimers. Each cDNA sample was run as two technical replicates and averaged for expression analysis. Samples were discarded for quantification if the difference between replicates was over 0,5 cycles.

To determine expression differences between CTRL and MCT8MO embryos, statistical analysis was carried out in GraphPad Prism v6.01 (San Diego, USA). The normality of the data was previously accessed using D’Agostino & Pearson omnibus normality test. Statistical significance was determined by unpaired Students t-test: two-sample, assuming equal variances (significance considered if p<0,05).

### Immunohistochemistry

One-cell stage embryos microinjected with either 0.8pmol of either CTRLMO (GeneTools) or MCT8MO ^13^ were fixed at selected stages in ice-cold 4%PFA/PBS overnight at 4°C. Samples were washed, depigmented when needed with PBS/0.3%H2O2/0.5%KOH, transferred into 100% methanol and stored at -20°C until use. Samples in 100% MeOH were brought to room temperature and washed using a MeOH:PBS series (100% MeOH to 100% PBS). Embryos were hydrated, washed in PBS with 0.1% Triton X-100 (PBTr) and blocked with the addition of 10% sheep serum (Sigma Aldrich). Primary antibodies used were: rabbit anti-HuC/D (1:500, 16A11 - Invitrogen), mouse anti-Zrf1 (1:100 - ZIRC) and anti-nkx6.1 (1:50; DSHB). Samples were washed, and secondary antibody fluorescent labelling was carried out using a goat anti-mouse IgG-CF594 (Sigma) or IgG-CF488 (Sigma) anti-serum (1/400). Imaging was carried out in a Zeiss Z1 light-sheet microscope. Images were imported into Fiji, and a region of interest was selected in a two-somite area (200μm) between somite 8-12. For neuron number determination, the 3D object counter in Fiji was used. The stained area was measured in Fiji for glial cell abundance after maximum intensity projection. The normality of the data was previously accessed using D’Agostino & Pearson omnibus normality test. Statistical significance was determined by unpaired Students t-test: two-sample, assuming equal variances (significance considered if p<0,05).

### Riboprobe preparation

To prepare *neurog1, fabp7a, slc1a2b, olig2* riboprobes for in situ hybridisation primers (Table 2) were designed using as template the sequences from the assembled transcriptome^15^. Following the manufacturer’s recommendation, an amplification and isolation of the cDNA of selected genes were carried out using a Thermo DreamTaq PCR kit. Amplified fragment isolated by agarose gel band extraction after electrophoresis and cloned into a pGemT easy vector described by the manufacturer (Promega, Germany). Isolated plasmid DNA was sequenced to confirm the identity and orientation of each clone.

**Table 2.**
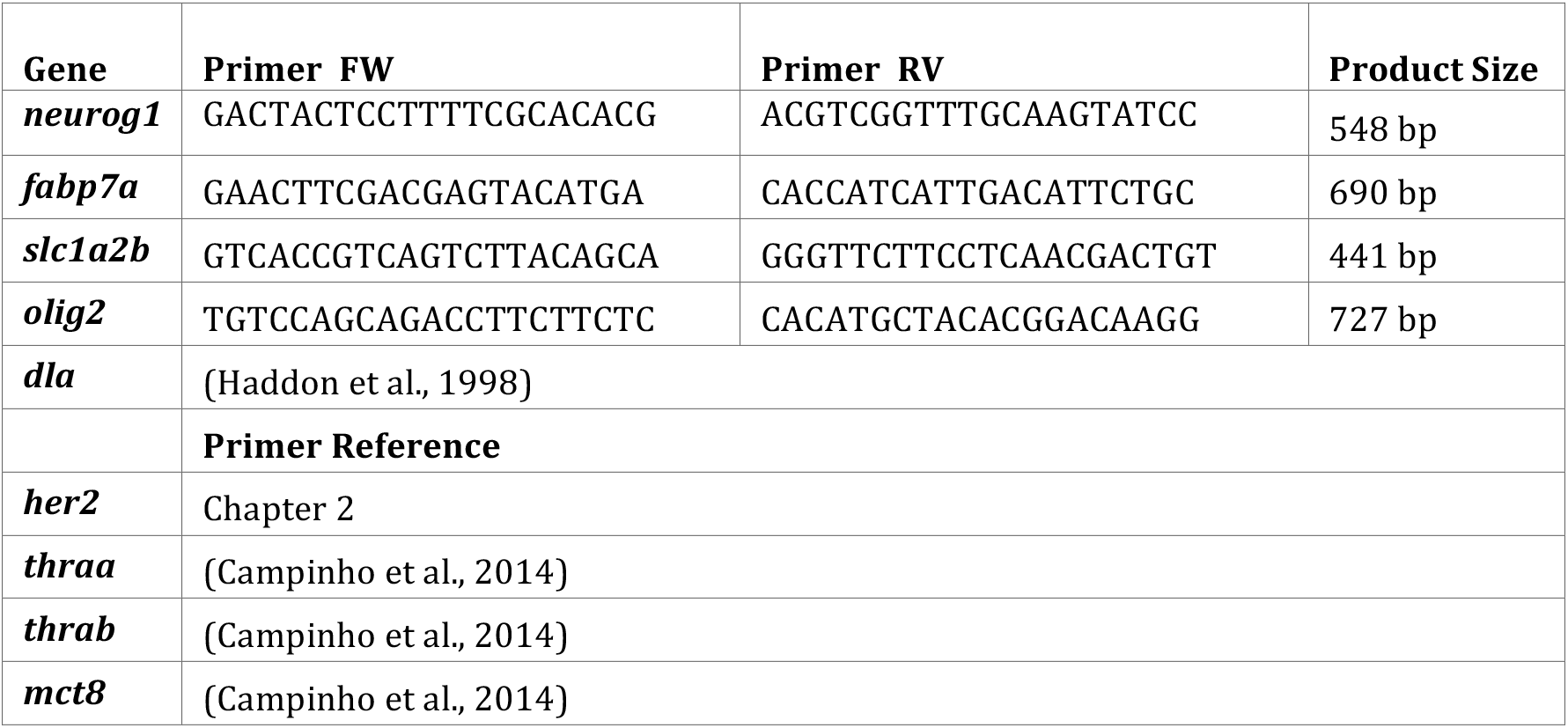
Primers used for cloning of target sequences, and references of other probes used.

The coding region of interest genes and the flanking T7 and SP6 RNA polymerase promoter were PCR amplified using M13 forward and reverse primers. PCR protocol was 95C 5minutes, 35 cycles of 95C for 30 seconds, 60C for 30 seconds and 1 minute at 72C, followed by a final extension set of 5 minutes at 72C. PCR products were purified by agarose gel and extracted using a GFX gel band extraction kit according to the manufacturer’s instruction (Omega Biotek, USA). 500ng of purified PCR fragments were used to prepare Digoxigenin or Fluorescein labelled antisense probes. These were synthesised by in vitro transcription with DIG-RNA labelling Kit or Fluorescein-RNA labelling kit (Roche, Switzerland). The integrity of probes was determined by gel electrophoresis and storage in 50% RNAlater (Sigma, USA) at -20°C.

### Whole-mount in-situ hybridisation (WISH)

Chromogenic WISH was carried out as previously described ^13^. Briefly, samples in 100% MeOH were brought to room temperature and washed using a MeOH:PBS series (100% MeOH to 0% MeOH) and finally rinsed several times in PBS/Tween-20 (PBSTw). Samples were digested with proteinase K (1 µg/mL) in 1×PBS from 5 to 20 minutes, depending on the embryonic stage. After permeabilisation, samples were re-fixed in 4% PFA/1×PBS and then pre-hybridised for 2 hours at 68 °C in a hybridisation mix (HybMix). HybMix was discarded and replaced by pre-warmed HybMix containing 0.25 ng/mL of Dig-labelled cRNA probe and hybridised overnight at 68 °C. Samples were then subject to stringency washes at 68 °C. Samples were incubated overnight at 4°C in 0.1M malic acid/0.1%-TritonX (MABTr)/10% sheep serum (Sigma-Aldrich)/2% Blocking solution (Roche, Switzerland) with anti-DIG-AP Fab fragments serum (1:5000, Roche, Switzerland). Samples were washed in MABTr and then incubated in NBT (nitro blue tetrazolium)/BCIP (bromo-chloro-indolyl-phosphate, Roche) for colour development, washed in stop solution. Zebrafish embryos (n≥10/stage/experimental condition) were transferred to 100% glycerol through a gradient and photographed under a stereoscope (Olympus) coupled to a digital colour camera (Optika).

To analyse the cell distribution pattern on transverse sections of the spinal cord, whole WISH individuals (*her2*, *fabp7a*, *neurog1* and *slc1a2b*) were re-fixed in PFA4%, dehydrated in MeOH/PBS and embedded in paraffin using isopropanol/paraffin gradient. Paraffin blocks were sectioned at 8µm and mounted on Poly-L-Lysine covered slides. Sections of interest were dewaxed, and coverslips mounted with glycerol-gelatine (Sigma). Images were photographed under a Leica LM2000 microscope coupled to a digital colour camera DS480 (Leica).

### Double fluorescent whole-mount in-situ hybridisation WISH

Riboprobes were generated and labelled with either digoxigenin (*mct8*, *thraa* and *thrab*) or fluorescein (*her2*, *dla* and *fabp7a*). The hybridisation with the probes was simultaneously, and antibody detection and development of the signal were carried out sequentially using a combination of antibody/Tyramide signal amplification (Perkin-Elmer, USA).

Briefly, samples in 100% MeOH were brought to room temperature and washed using a MeOH:PBS series (100% MeOH to 0% MeOH) and finally rinsed several times in PBS/Tween-20 (PBSTw). Samples were digested with proteinase K (1 µg/mL) in 1×PBS from 5 to 20 minutes, depending on the embryonic stage. After permeabilisation, samples were re-fixed in 4% PFA/1×PBS and afterwards pre-hybridised for 2 hours at 68 °C in HybMix. HybMix was discarded and replaced by pre-warmed HybMix containing 0.5 ng/mL of Dig- and Fluorescein-labelled cRNA probes and hybridised overnight at 68 °C. Samples were then subject to stringency washes at 68 °C. For first probe detection, embryos were incubated overnight at 4 °C in blocking solution MABTr/10% sheep serum (Sigma-Aldrich)/2% Blocking solution (Roche, Switzerland) with anti-DIG-POD Fab fragments serum (1:500, Roche, Switzerland). Embryos were washed in PBSTw and then incubated in Alexa Fluor-594 Tyramide Reagent (ThermoFisher, USA), 1:100 in amplification reagent (Perkin Elmer) for fluorescent colour development, according to the instructions, followed by several washes in PBSTw. For second probe detection, peroxidase activity of previous POD conjugated anti-serum was quenched by incubating samples for 1h in 3% H2O2 in PBS. Samples were washed in PBSTr and incubated overnight at 4 °C MABTr/10% sheep serum (Sigma-Aldrich)/2% Blocking solution (Roche, Switzerland) with anti-Fluorescein-POD Fab fragments serum (1:500, Roche, Switzerland). Embryos were washed in PBSTw and then incubated in FITC-Tyramide (Perkin-Elmer) 1:100 in amplification reagent (Perkin Elmer), followed by several washes in PBSTw. Samples were stored in PBS containing 0,1% Dabco (CarlRoth, Germany). Fluorescent imaging was carried out immediately to avoid bleaching of the fluorophores, using a Lightsheet Z.1 (ZEISS) microscope. Samples were mounted in 1% low melting agarose (CarlRoth, Germany) and imaged using dual illumination and a z step of 1,69µm or 1,813 µm depending on the zoom used. The total depth of the medial spinal cord was acquired using a 10x lens with 2.5x or 1x optical zoom. Briefly, dual illumination image volumes from the Z.1 were merged using dual side fusion option (Zen Black, Zeiss) images.

For imaging and co-localisation analysis, acquired images were imported into Fiji ^58^, ROI was selected in a two somite area (200μm) between somite 8-13. The threshold was adjusted and fixed for each gene pair. The Fiji Colocalization Colormap plugin ^59^ was used to determine co-localisation. At 3-8 individuals per condition were analysed. The resulting stack of the co-localisation was then superimposed into the original Z1 image to create the final figures, and when necessary, stacks were resliced in y to enable lateral views.

### Mitosis detection

Immediately after fluorescent *wish*, embryos were subject to immunohistochemistry to detect mitotic cells. The primary antibody used was rabbit anti-PH3 1:500 (Millipore), and the secondary antibody goat anti-rabbit IgG-CF594 (Sigma). Antibody incubation and blocking steps were performed in 1xPBS:10%Sheep serum.

### Apoptosis detection

Immediately after fluorescent *wish*, embryos were washed for 15 minutes at RT with 1xPBS/0.1% TritonX 100 (Sigma)/0.1 M Sodium Acetate pH6. Embryos were further treated 15 minutes at RT with 1ug/mL Proteinase K (Sigma) followed by four 5 minutes washes in 1xPBT. According to the manufacturer’s instructions, including experimental controls, apoptotic cells in experimental animals were labelled using the Roche in situ cell death TMR-red detection kit.

### Imaging of *wish*-apoptosis and *wish*-mitosis

Lightsheet Z.1 (ZEISS) microscope was used to acquire images of *wish*-apoptosis *and wish*-mitosis. Samples were mounted in 1% low melting agarose (CarlRoth, Germany) and imaged using dual illumination and a z step of 1,69µm or 1,813 µm depending on the zoom used. The total depth of the medial spinal cord was acquired using a 10x lens, 2.5x or 1x zoom. Briefly, dual illumination images from the Z.1 were merged using Dual side Fusion (Zen Black, Zeiss) images were then imported into Fiji, ROI was selected in a two somite area (200μm) between somite 9-12. The threshold was adjusted and fixed for each target, and the Colocalization Colormap plugin ^59^ was used to determine co-localisation. Co-localized cells were counted manually with Fiji’s "3D object counter" tool.

### NICD Overexpression

pCS2+ Plasmids containing the cDNA coding for CAAX-GFP (membrane label) and the Notch-intracellular domain (NICD) ^60^ were linearised, and mRNAs synthesised using the mMessage Machine SP6 transcription kit from Ambion, following the manufacturer’s instructions. The mRNAs were phenol: chloroform purified, diluted in RNAse free water and frozen at -80C until use. In this experiment, a variant of zebrafish notch1a, notch1a-intracellular domain (NICD), was used, which encodes a Notch receptor that is constitutively active in neurogenesis. The effects of this NICD mRNA injection are attributed to high NOTCH activity in general (Takke & Campos-Ortega 1999).

### Live Imaging

Zebrafish Tg(*elav3:LY-mCherry*)^61^ x WT AB previously injected with CTRL or MCT8MO were used for mRNA injection. For mosaic overexpression of NICD, 100 pg of NICD mRNA and/or 50pg of GFP-CaaX mRNA were injected into one blastomere (DRA1, DRA2) between the 16- to 32-cell stages. GFP-CaaX mRNA was used to allow the individual cell visualisation of the cell-autonomous response to NICD in CTRL and MCT8MO injected embryos. Hence, four groups were considered, GFP-CaaX injection in MCT8MO and CTRLMO; NICD and GFP-CaaX injection in MCT8MO and CTRLMO. Embryos were left to develop at 28°C until 22hpf when sorting and mounting for imaging was performed.

Imaging was carried out by light-sheet microscopy, Lightsheet Z.1 (ZEISS, Germany), as described previously^62^, with minor alterations. Briefly, embryos were anaesthetised with 0,08% Tricane pH7.4 buffered, mounted alive in 0.3% (w/v) low-melting agarose (LMA) in E3 medium containing tricaine (0.08%) into FEP tubes closed with a 1% LMA. Three animals per group CTRLMO and MCT8MO were imaged in the same tube. Two independent experiments were carried out. Time lapses images were taken from 23 until 26hpf. Z-stacks ranging the full depth of the medial spinal cord were acquired every 15 min during 3h. The spinal cord was imaged with x20 lens, 2x zoom with a z-step of 1.56 μm with single angle and dual illumination.

For image analysis, dual illumination images from the Z.1 were merged using Dual side Fusion (Zen Black, Zeiss). Images were imported into Fiji, and a region of interest was selected in a two somite area (200μm) between somite 8-12. Analysis of cell divisions was performed manually in FIJI. Only Huc(-) cells expressing GFP were tracked for analysis.

Statistical analysis was done using Graphpad Prism v6.01 (San Diego, USA). Values are represented as means ± SD. The normality of the data was previously accessed using D’Agostino & Pearson omnibus normality test. Symmetric divisions of GFP+ cells in any group were considered if the cell division plane was 0-<30° to the ventricular side of the spinal cord. Asymmetric divisions were considered if the cell division plane was ≥30-90° to the ventricular side of the spinal cord ^23, 54^. Distribution differences in symmetric/asymmetric divisions between experimental groups were determined after X2 analysis. Statistical difference in symmetric or asymmetric divisions between experimental groups was determined by one-way ANOVA followed by Holm-Sidak’s multiple comparison post hoc analysis. Significance was considered if p<0,05.

## Acknowledgments

The authors wish to thank Susana Santos Lopes for the Notch-intracellular domain (NICD) construct and Michael Orger for the zebrafish Tg(*elav3:LY-mCherry*) line.

This study received Portuguese national funds from FCT - Foundation for Science and Technology through project project PTDC/EXPL/MAR-BIO/0430/2013 and FCT UIDB/04326/2020. COMPETE 2020 through project EMBRC.PT ALG-01-0145-FEDER-022121. ABC-RI CRESC Algarve 2020. NS is a recipient of a FCT PhD grants SFRH/BD/111226/2015. MAC is a recipient of a FCT-IF Starting Grant (IF/01274/2014).

## Supplementary Figure Legends

**Supplementary Figure 1.**
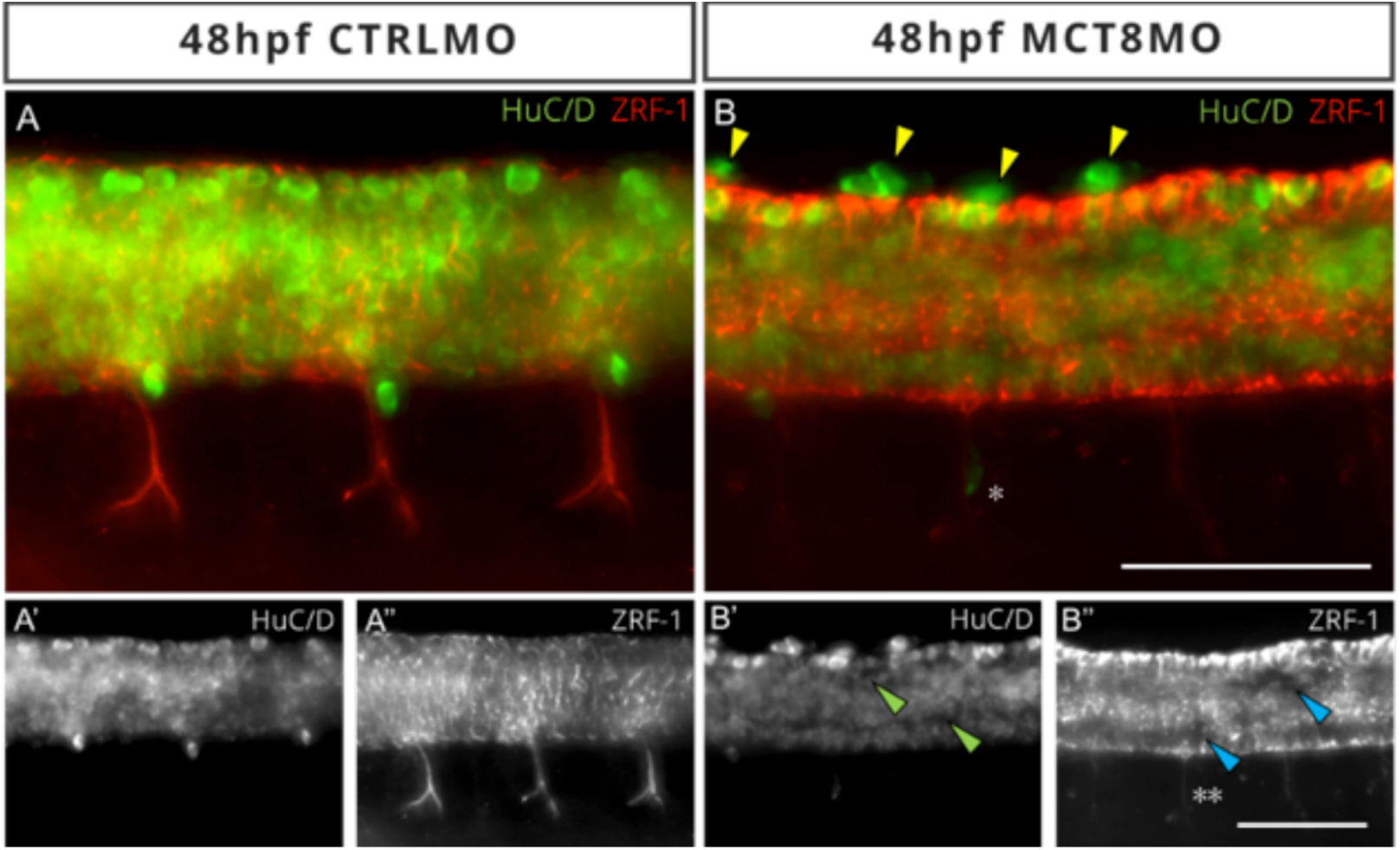
At the end of embryogenesis (48hpf) the cytoarchitecture of the spinal cord is altered in embryos with impaired MT3 signalling. Characterization of the spinal cord of 48hpf zebrafish by double immunohistochemistry labelling of mature neurons (HUC/D) and glial cells (ZRF-1). (A) In control morpholino embryos (CTRLMO), neurons are distributed in a V-shaped organization (A’), and the perineurial glial cells (red) are located adjacent to the motor nerve, descending toward the muscle. Glial cells are well distributed and surround neurons in all directions(A’’). (B) In MCT8 morphant embryos (MCT8MO), neurons accumulate ectopically outside the dorsal limit of the spinal cord (yellow arrowheads in B) and are not distribuited equally along the spinal cord (green arrowheads in B’). Perineurial glial cells projections are absent or abnormally extended to the myotome (**). Glial cells are disorganized and have accumulated in the most dorsal and ventral regions leading to "holes" in the spinal cord (blue arrowheads in B’’). Motoneurons do not migrate normally towards the myotome in MCT8MO embryos (* in B). In all images dorsal up, anterior left. Images are maximum projections of spinal cord z-stacks. Scale bar represents 50 µm.

**Supplementary Figure 2.**
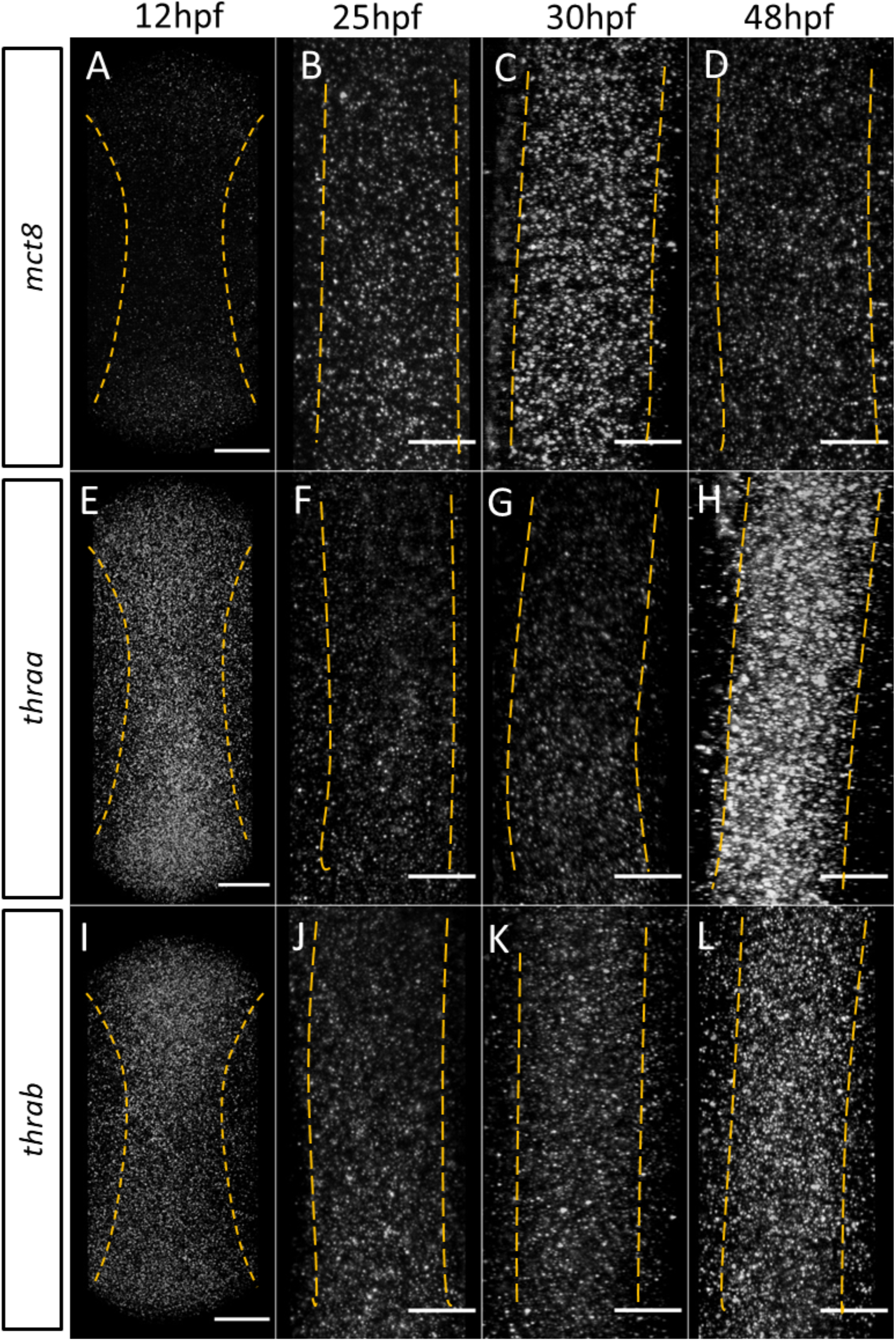
Spatial temporal expression of MT3 cellular signaling genes mct8, thraa and thrab during zebrafish spinal cord neurodevelopment. Fluorescent WMISH in CTRLMO embryos allowed determination of the spatio-temporal expression pattern of *mct8* (A-D), *thraa* (E-H)*, and thrab* (I-L) at 12 (A, E and I), 25 (B, F and J), 30 (C, G and K) and 48hpf (D, H and L). Image represent dorsal maximum projections of the whole spinal cord at 12hpf (A-I) and section of the spinal cord between somites 8-12 at 25 (B, F and J), 30 (C, G and K) and 48hpf (D, H and L). Yellow *v*ertical dashed lines show the lateral boundary of the spinal cord. In all images the anterior is upwards. Scale bars represent 25µm, except for 12hpf where the scale bar represents 50 µm.

**Supplementary Figure 3.**
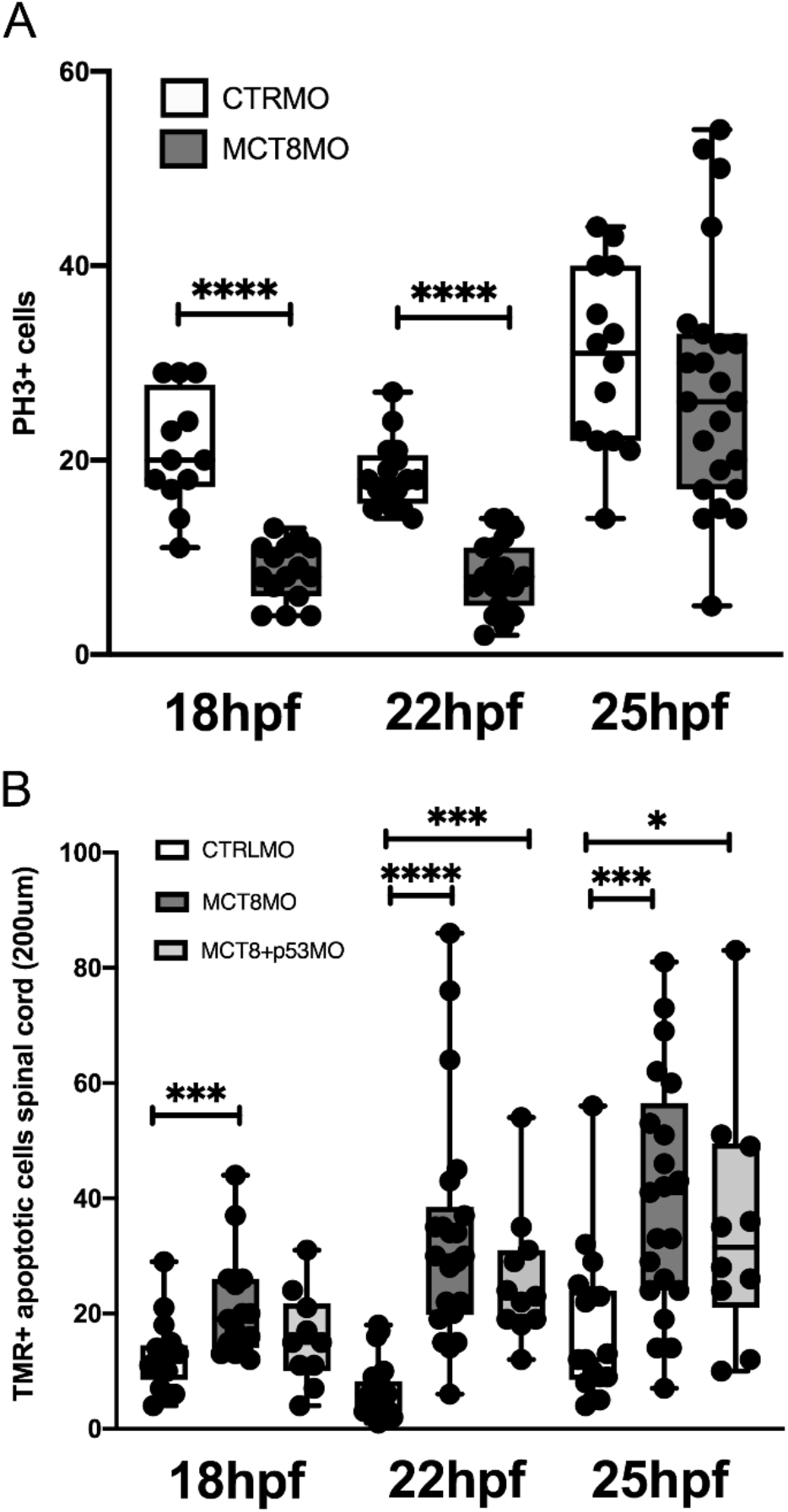
Impaired MT3 signalling affects both mitosis and apoptosis in the spinal. Overall mitotic (A) and apoptotic (B) cells in spinal cord of embryos injected with control (CTRLMO) or mct8 morpholino (MCT8MO) at 18, 22 and 25hpf. Quantifications were carried out in a 200μm wide volume of the spinal cord between somite 8-12. In the case of apoptosis it was also used an additional experimental group where it was co-injected with the MCT8MO a p53MO to mitigate effects in apoptosis due to non-specific toxicity effects of MCT8MO. No differences were found between MCT8MO and MCT8+p53MO group. Statistical differences were found after unpaired t-test and considered if p≤0.05. * p≤0.05, *** p<0.001, **** p<0.0001.

